## Supplementary Information for "HEXIM1 inter-monomer autoinhibition governs 7SK RNA binding specificity and P-TEFb inactivation"

**List of supplementary figures and table:**

**Supplementary Fig. 1: Sequence alignment of Hexim1 and Hexim2 orthologs displaying the N-terminal variable region.**
**Supplementary Fig. 2: Sequence alignment of Hexim1 and Hexim2 orthologs displaying the conserved regions.**
**Supplementary Fig. 3: Prediction of Hexim1 disordered regions by a language model DR-BERT.**
**Supplementary Fig. 4: Native PAGE and mass photometry studies of Hexim1 monomeric and dimeric proteins.**
**Supplementary Fig. 5: NMR assignments of Hexim1 BR-L-AR.**
**Supplementary Fig. 6: NMR binding studies of SL1-dI RNA with Hexim1 BR-L-AR.**
**Supplementary Fig. 7: $^{13}$C-$^{1}$H HSQC assignments of SL1-dI RNA.**
**Supplementary Fig. 8: NMR binding studies of SL1-dI$_{\Delta U}$ RNA with Hexim1 BR-L-AR.**
**Supplementary Fig. 9: NMR titration of SL1-dI into Hexim1 BR-L-AR.**
**Supplementary Fig. 10: Spectral behavior of Hexim1 BR-L, BR and BR$_{ARM1}$ upon binding 7SK RNA.**
**Notes about line broadening due to RNA binding**
**Supplementary Fig. 11: $^{15}$N BR-L-AR(S237C$^{MTSL}$) broadens more severely than S226C$^{MTSL}$ throughout the protein sequence.**
**Supplementary Fig. 12: $^{15}$N-$^{1}$H HSQC spectral overlay of $^{15}$N BR-L-AR(A210C)**
**Supplementary Fig. 13: $^{15}$N-$^{1}$H HSQC spectral overlay of $^{15}$N BR-L-AR(S183C)**
**Supplementary Fig. 14: Binding studies of SL1 RNA with Hexim1 by EMSA.**
**Notes for EMSA binding curve fittings**
**Supplementary Fig. 15: Fitting results from EMSA studies.**
**Supplementary Fig. 16: NMR binding studies of SL1-dII RNA with Hexim1 BR-L.**
**Supplementary Fig. 17: NMR binding studies of SL1-d RNA with Hexim1 BR$_{ARM1}$.**
**Supplementary Fig. 18: NMR binding studies of SL1-dIIm RNA with Hexim1 BR$_{ARM1}$.**
**Supplementary Fig. 19: NMR binding studies of SL1-pII RNA with Hexim1 BR$_{ARM1}$.**
**Supplementary Fig. 20: NMR binding studies of SL1-p RNA with Hexim1 BR$_{ARM1}$.**
**Supplementary Fig. 21: NMR binding studies of SL1-mp RNA with Hexim1 BR$_{ARM1}$.**
**Supplementary Fig. 22: NMR assignments and binding study of circular 7SK SL1alt RNA.**
**Supplementary Fig. 23: Longer incubation time reveals secondary phosphorylation sites in αL helix.**
**Supplementary Fig. 24: ITC measurements of NBLA and Hexim1 dimer binding to 7SK RNA constructs.**
**Supplementary Fig. 25: Hexim1 dimer variants that disrupt autoinhibition became non-specific RNA binder.**
**Supplementary Fig. 26: Sequence alignment of HIV-1 Tat protein.**
**Supplementary discussion regarding N-terminus of Hexim and Tat.**
**Supplementary Table 1: Thermodynamic parameters for binding of Hexim1 to 7SK RNA from ITC with replicate numbers**
**Uncropped gel images.**

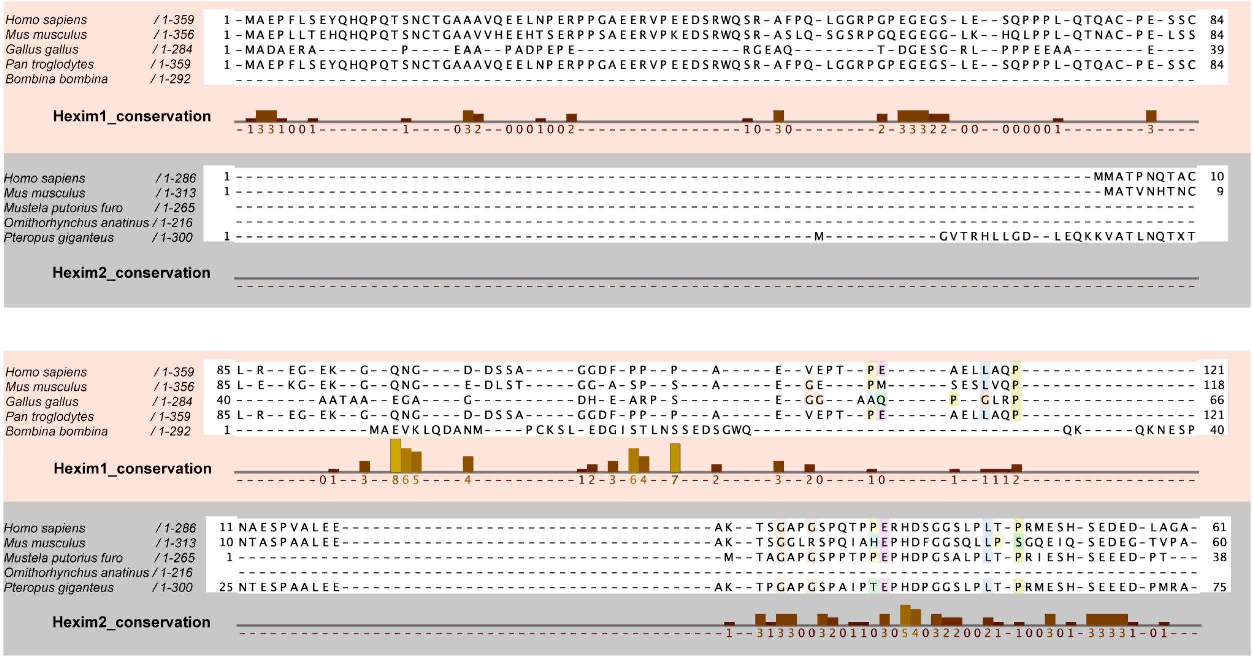

**Supplementary Fig. 1: Sequence alignment of Hexim1 and Hexim2 orthologs displaying the N-terminal variable region.** A total number of 294 genes for Hexim1 and 210 genes for Hexim2 were aligned. Three representative Hexim1 and Hexim2 genes each, in addition to human and mouse (making five total), are randomly selected for this display, shown in red and grey boxes, respectively. Hexim1 orthologs have longer N-terminal region than Hexim2. Empty columns mostly reflect distinction between Hexim1 and Hexim2 genes. The full alignment from the total gene pools showed significantly more variations and insertions than these five representative genes used for display, which additionally informs on the poor conservation of N-terminal region.

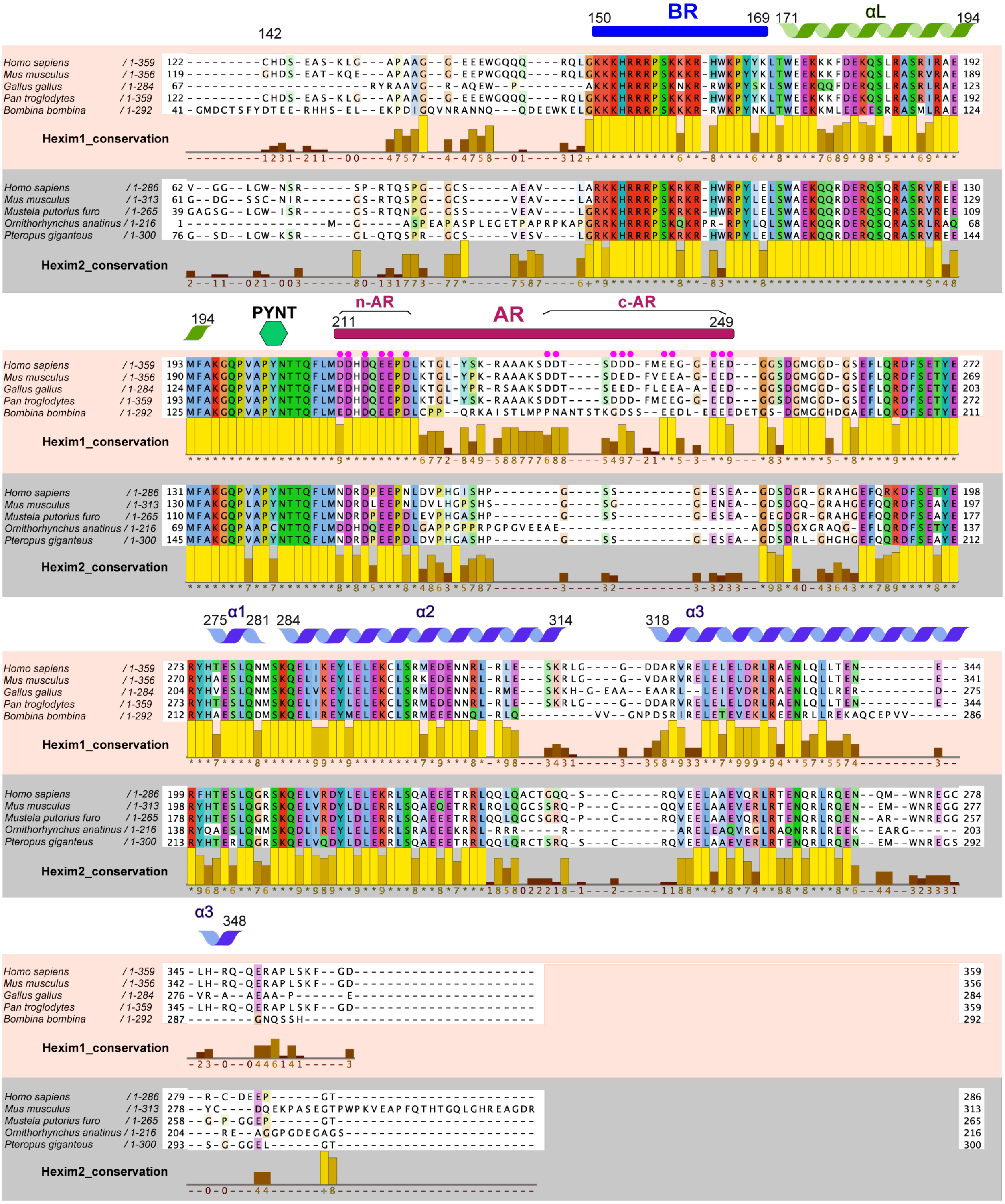

**Supplementary Fig. 2: Sequence alignment of Hexim1 and Hexim2 orthologs displaying the conserved regions: BR, αL, PYNT motif, AR and CC.** A total of 294 genes for Hexim1 and 210 genes for Hexim2 were aligned. Three representative Hexim1 and Hexim2 genes each, in addition to human and mouse, were randomly selected for this display, shown in red and grey boxes, respectively. The conservation scores are for the five selected homologs. Phe240 and Met241 are highly conserved with scores of 10 (+) and 8 in the full sequence alignment of 294 Hexim1 genes, of which the scores are much lower here due to the *Bb* gene having more sequence variation in this region. Note that Hexim2 orthologs lack the C-terminal acidic patch (c-AR), which is a conserved feature in Hexim1 orthologs. The coiled-coil domain (CC) comprises α1, α2 and α3 and a functionally indispensable flexible hinge between α2 and α3[1].

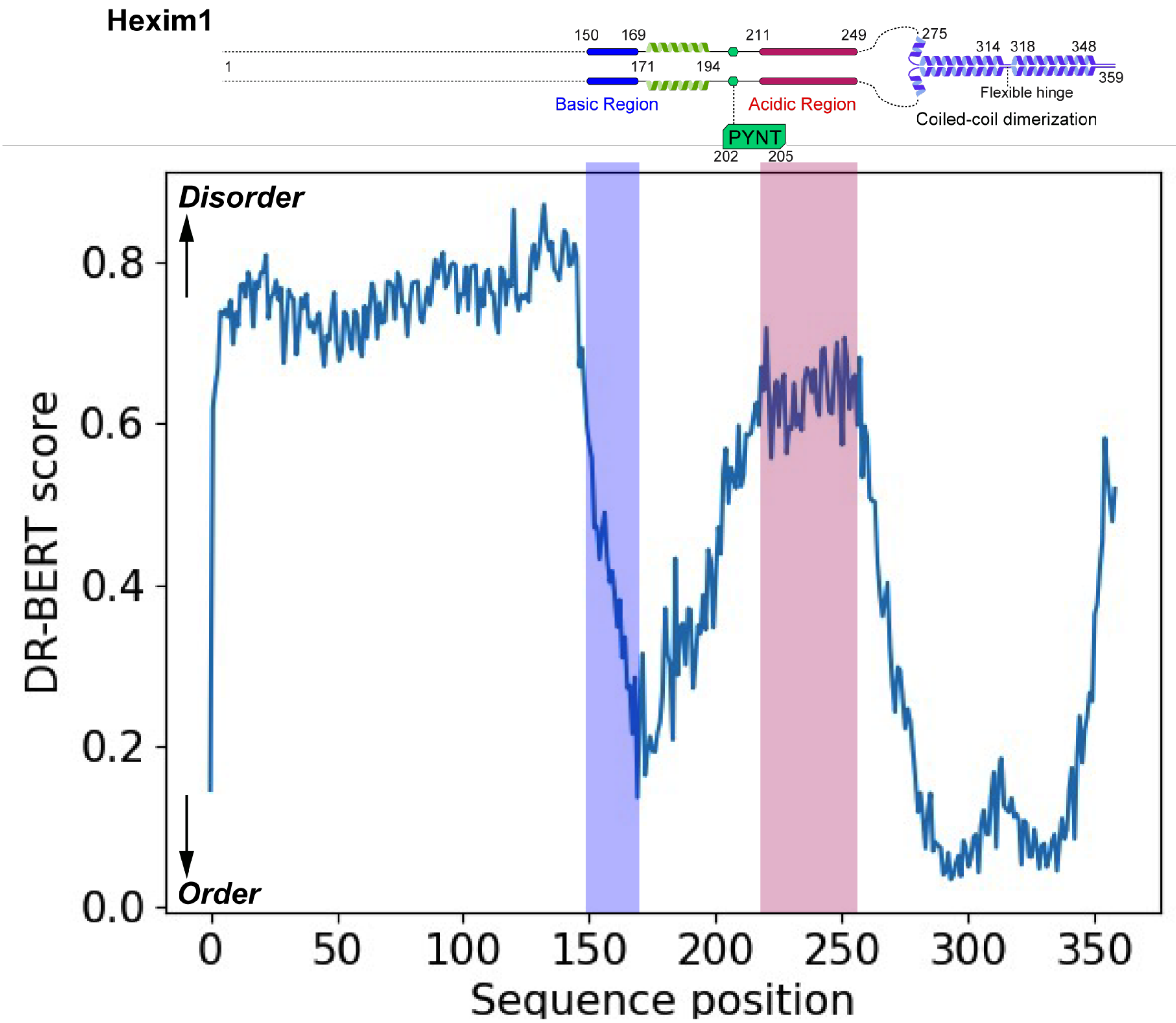


**Supplementary Fig. 3: Prediction of Hexim1 disordered regions by a language model DR-BERT[2].** Domain diagram of Hexim1 homodimer is shown at top.

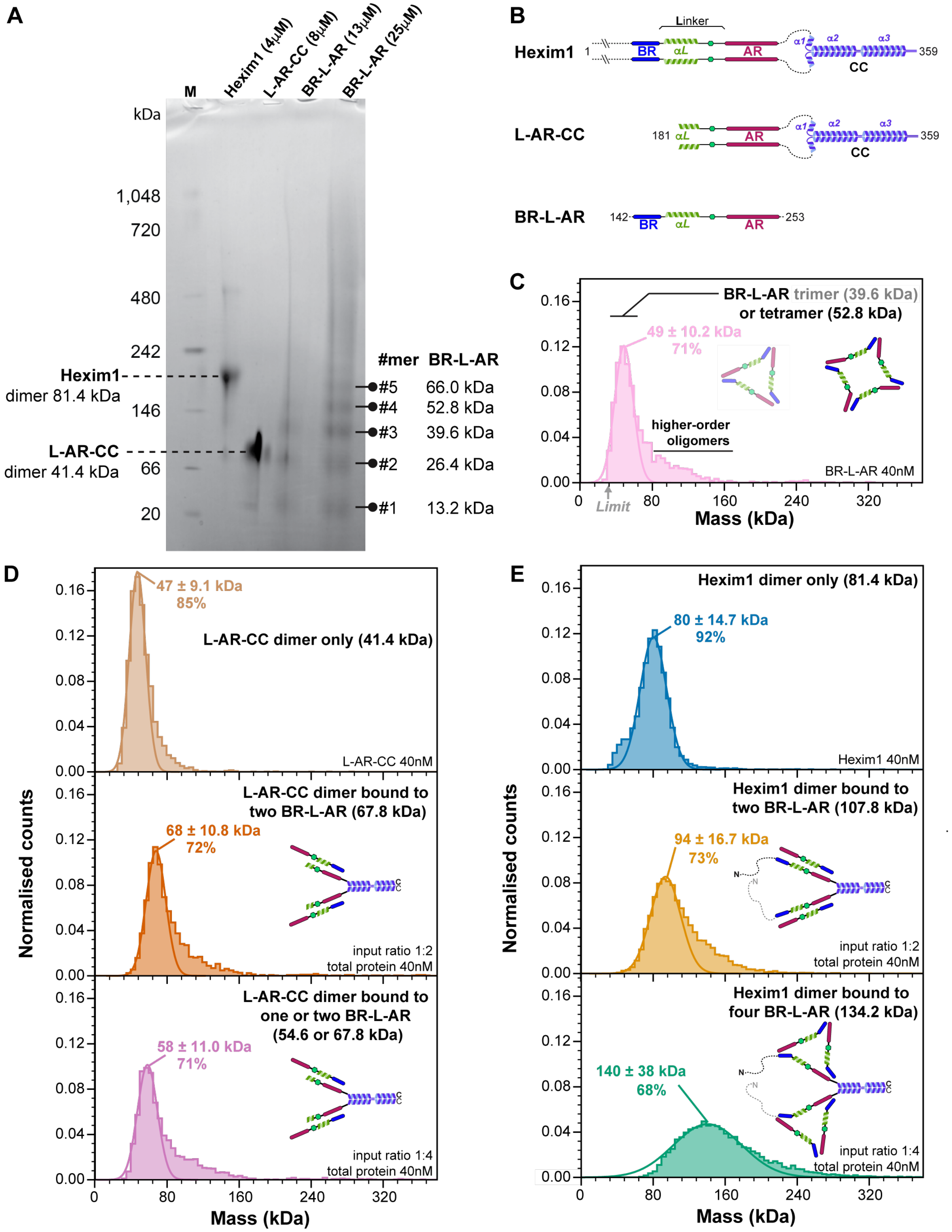


**Supplementary Fig. 4: Native PAGE and mass photometry studies of Hexim1 monomeric and dimeric proteins show stable intermolecular interactions under dilute conditions. A** Native PAGE analysis of freshly purified Hexim1 dimer, L-AR-CC dimer, and monomeric BR-L-AR protein constructs. Concentrations of proteins loaded in the lanes are labeled on top of the gel. Theoretical molecular weights of the three protein constructs are indicated on the two sides of the gel. Note that a small amount of tetramer (~6.7%) can be observed for Hexim1 at 4μM concentration, but not detected at 40nM concentration by Mass Photometry (panel E). **B** Domain diagrams of these three protein constructs are shown. **C** Mass photometry result of BR-L-AR detected existence of trimer and tetramer populations. The observed Mass Photometry peak corresponds to a molecular weight of 49 ± 10.2 kDa—between the theoretical trimer (39.6 kDa) and tetramer (52.8 kDa) sizes, but closer to the latter. This shift likely results from the trimer peak being near the 30 kDa detection limit (gray arrow), causing partial peak truncation and overlap with the tetramer peak. Monomer and dimer species fall below the detection threshold and are not observed. The percentages are derived from the Mass Photometry analysis, which calculates the proportion of total particle counts represented by the Gaussian-fitted peaks. **D** Mass photometry result of L-AR-CC dimer indicates homogeneous dimeric particles. In the presence of BR-L-AR, a stable 1:2 homodimer of heterodimers between L-AR-CC and BR-L-AR forms, which persists with a 1:4 input protein ratio due to L-AR-CC construct lacking BR. **E** Mass photometry result of Hexim1 dimer indicates homogeneous dimeric particles. In the presence of BR-L-AR, a stable homodimer of heterodimers between Hexim1 and BR-L-AR forms at 1:2 (1 Hexim1 dimer with 2 BR-L-AR monomers) input protein molar ratio, and a stable homodimer of heterotrimers between Hexim1 and BR-L-AR forms at 1:4 (1 Hexim1 dimer with 4 BR-L-AR monomers) input protein molar ratio. The cartoon illustrates the mode of intermolecular interactions, which is consistent with the trimer species detected in panels A and C.

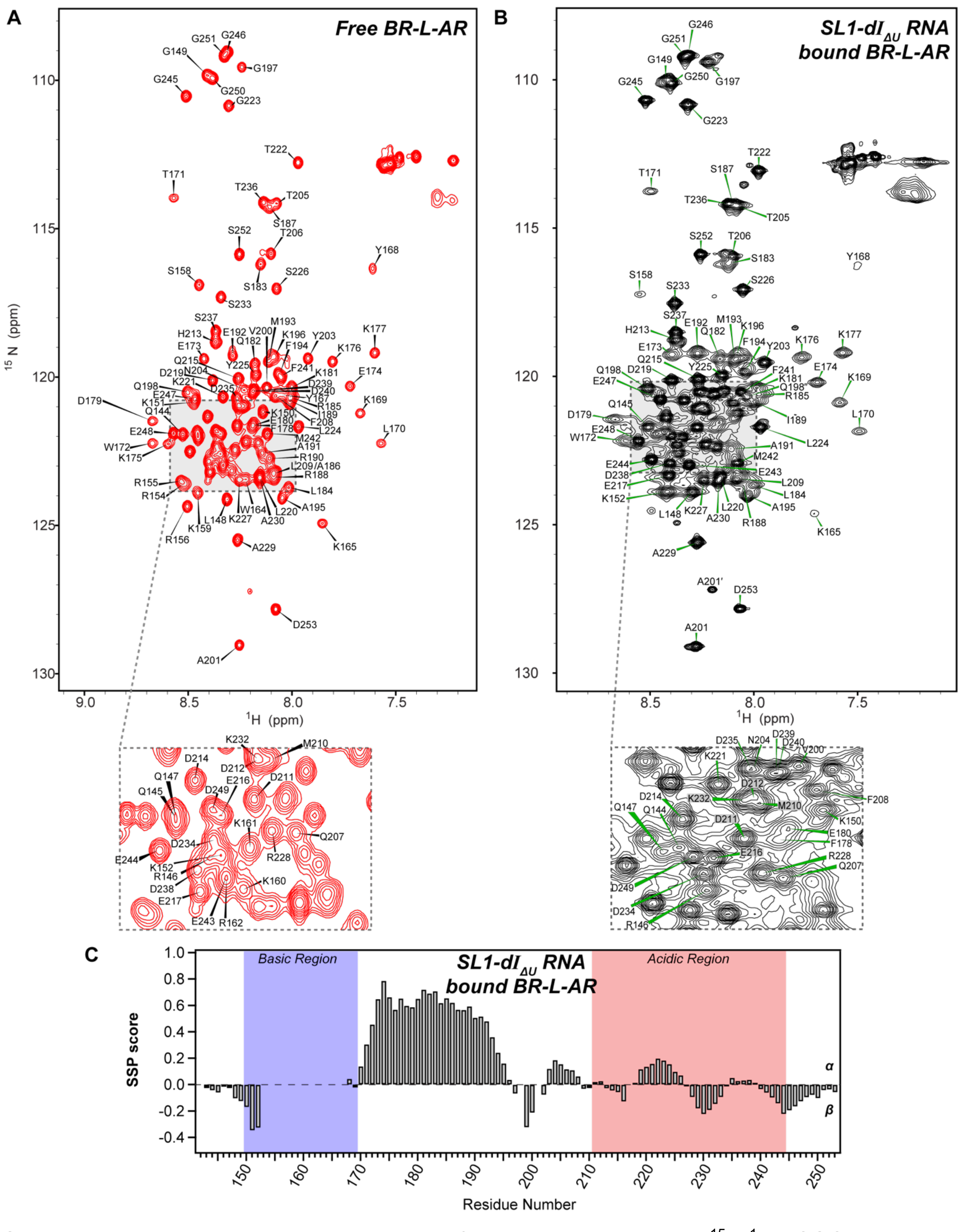


**Supplementary Fig. 5: NMR assignments of Hexim1 BR-L-AR. A-B** $^{15}$N-$^{1}$H HSQC spectra with labeled amide assignments in the absence (A) and presence (B) of SL1-dI$_{\Delta U}$ RNA at 600MHz. **C** Secondary structure propensity (SSP) scores of SL1-dI$_{\Delta U}$ RNA-bound BR-L-AR.

Note that the SSP scores of S158 and K165 that were assigned amide resonance in Fig. 2F are lacking values here, due to the lack of Cα/Cβ assignments for S158 in the RNA-bound form, and lack of Cβ chemical shift for K165.

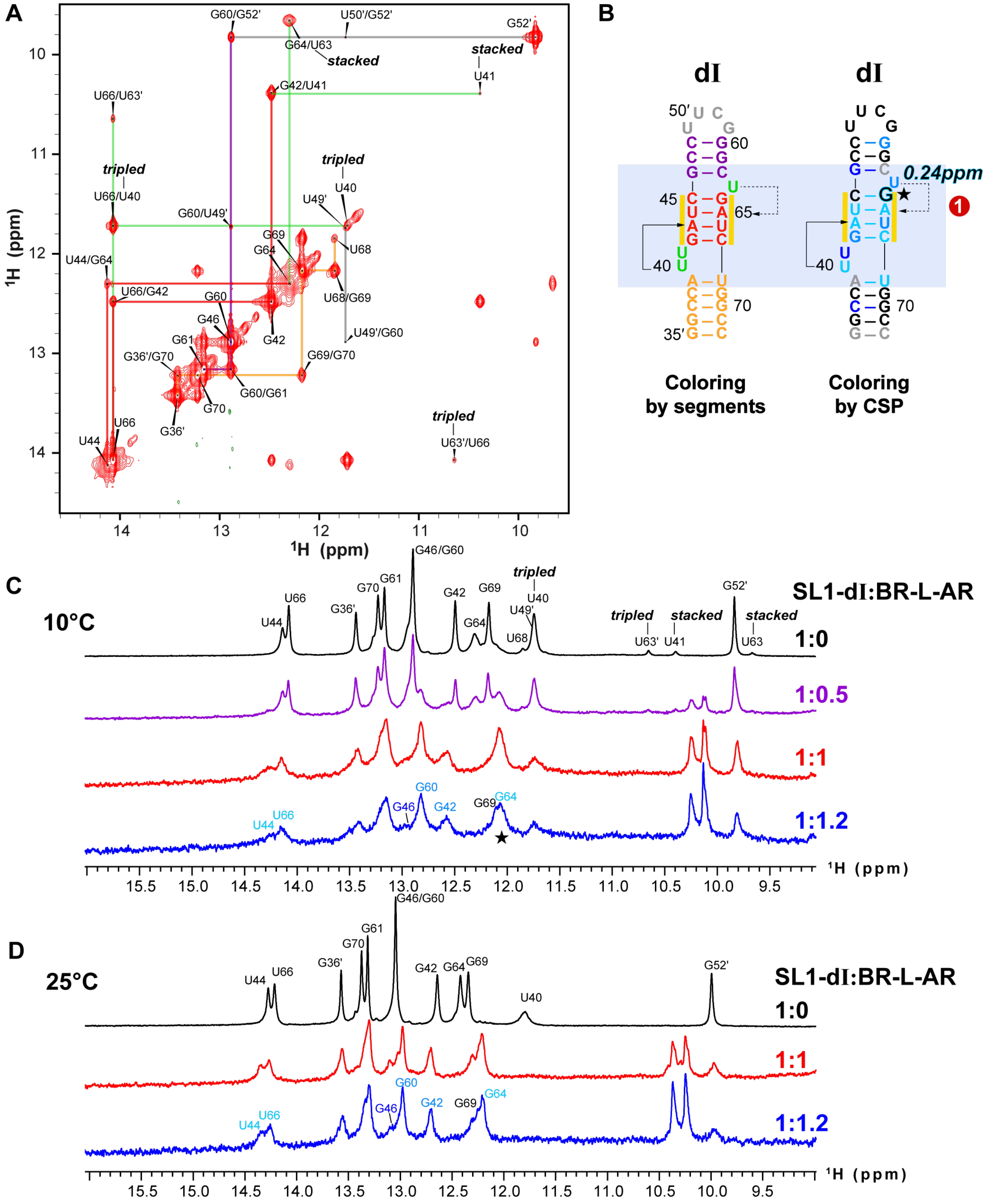


**Supplementary Fig. 6: NMR binding studies of SL1-dI RNA with Hexim1 BR-L-AR. A** 2D imino NOESY assignments of SL1-dI RNA. **B** Secondary structure of SL1-dI RNA colored by segments and by Chemical shift perturbations (CSP) upon binding BR-L-AR, respectively. The nucleotide experiencing the largest shift is highlighted with black/cyan font and a star, with the corresponding CSP value labeled. Severely broadened and highly overlapping resonances were excluded from the free/bound pair for CSP calculations. **C** 1D imino overlay of SL1-dI in the absence and presence of BR-L-AR at 10°C. Note that CSPs were calculated from all assigned protons from pairs of free and bound 2D spectra, instead of just 1D imino displayed here. **D** 1D imino overlay of SL1-dI in the absence and presence of BR-L-AR at 25°C.

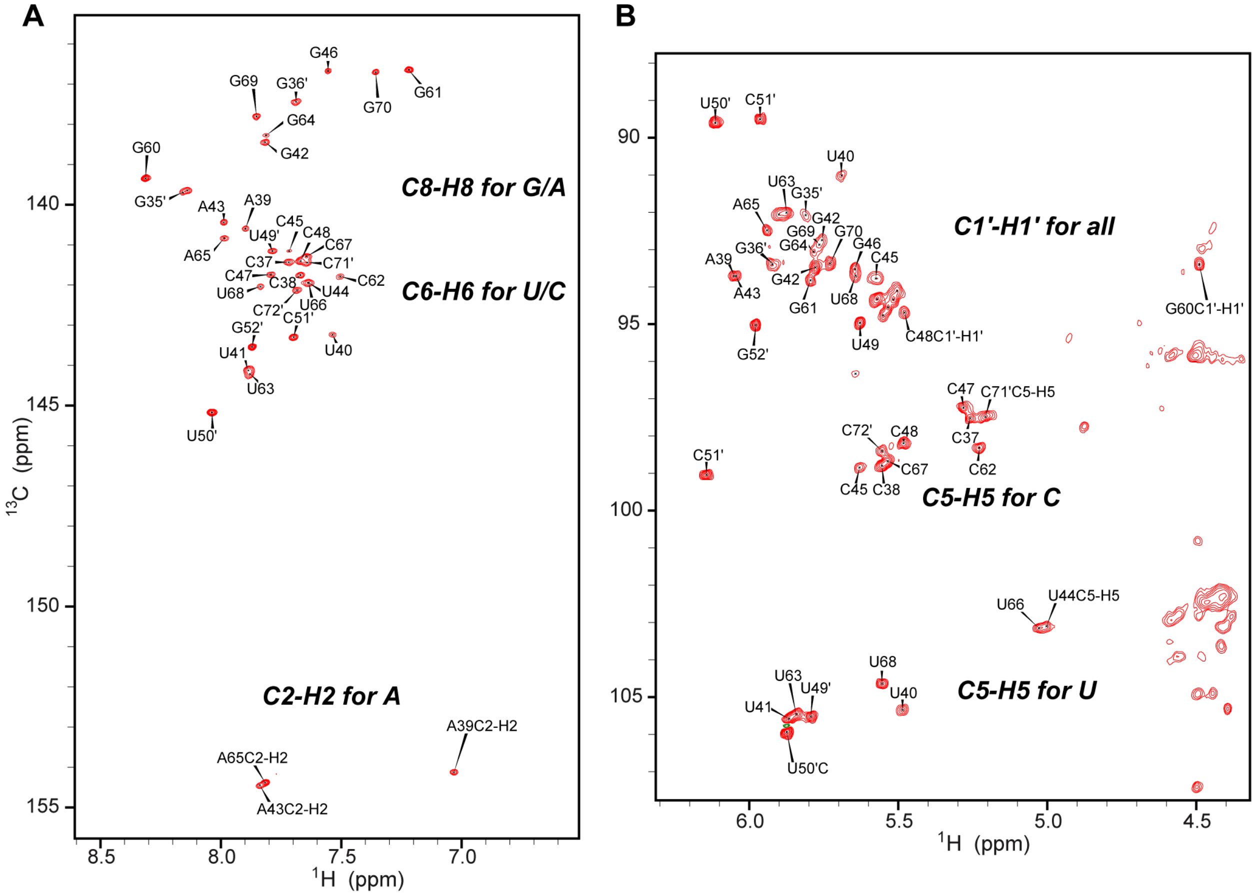


**Supplementary Fig. 7: $^{13}$C-$^{1}$H HSQC assignments of SL1-dl RNA. A** Aromatic $^{13}$C-$^{1}$H HSQC with assignments of C8-H8 for G and A, C6-H6 for U and C, as well as C2-H2 for A. **B** $^{13}$C-$^{1}$H HSQC with assignments of C1´-H1´ for all nucleotides, and C5-H5 for U and C.

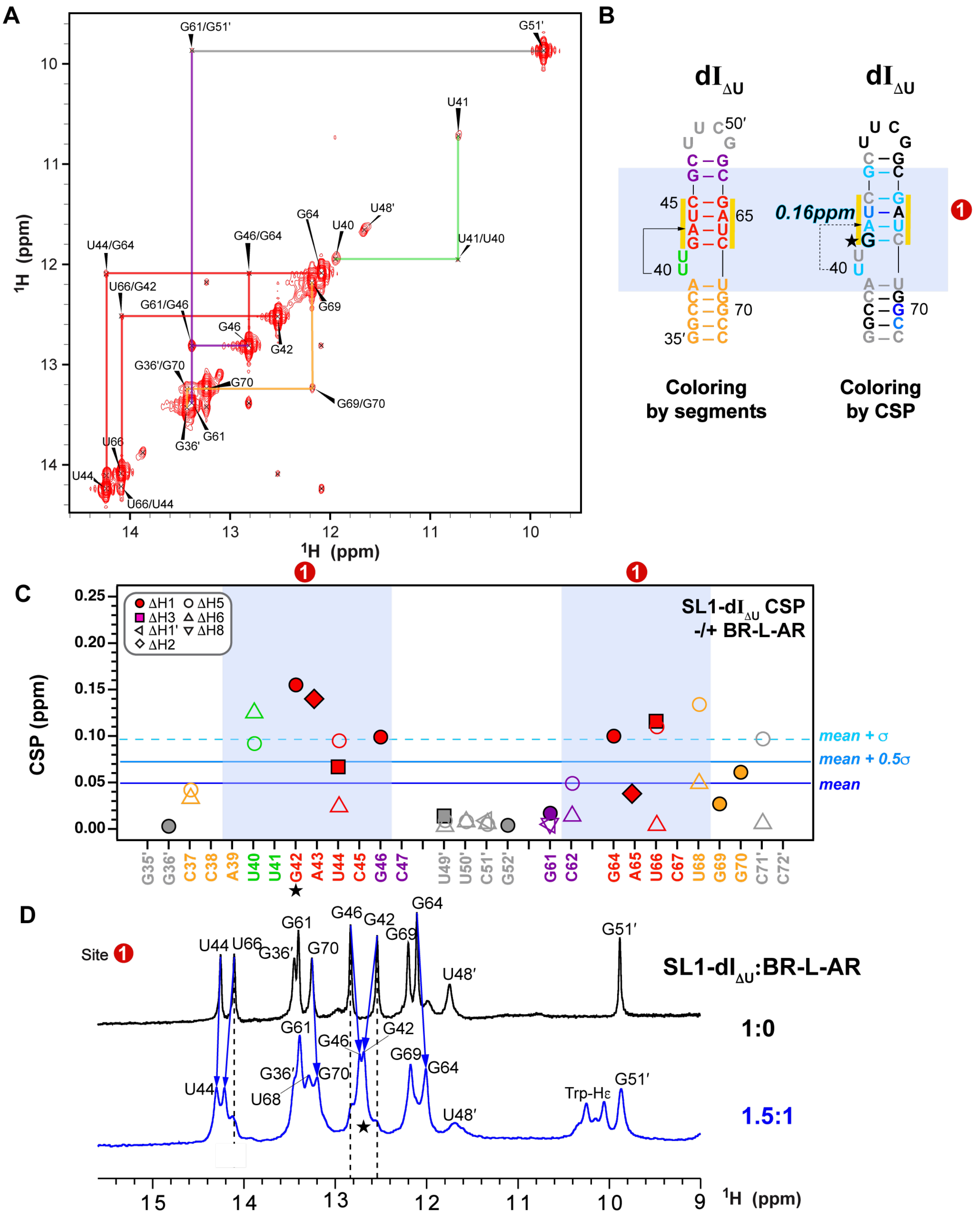


**Supplementary Fig. 8: NMR binding studies of SL1-dI$_{\Delta U}$ RNA with Hexim1 BR-L-AR. A** 2D imino NOESY assignments of SL1-dI$_{\Delta U}$ RNA. **B** Secondary structure of SL1-dI$_{\Delta U}$ RNA colored by segments and by Chemical shift perturbations (CSP) upon binding BR-L-AR, respectively. The nucleotide experiencing the largest shift is highlighted with black/cyan font and a star, with the corresponding CSP value labeled. **C** CSP plot of SL1-dI$_{\Delta U}$ RNA upon binding BR-L-AR. Severely broadened and highly overlapping resonances were excluded from the free/bound pair for CSP calculations. **D** Proton 1D imino region of linear 7SK SL1-dI$_{\Delta U}$ in the absence and presence of BR-L-AR at 600MHz.

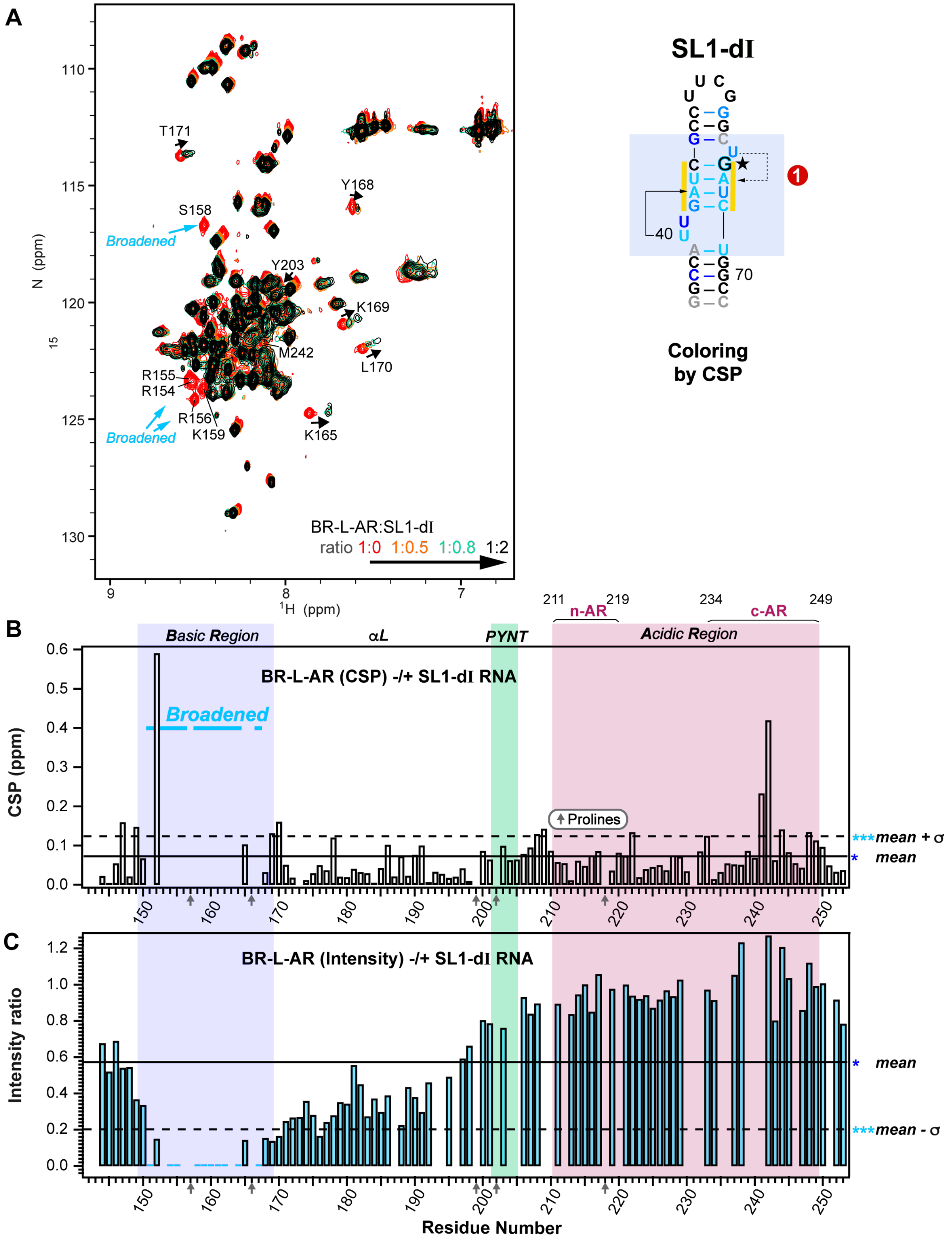


**Supplementary Fig. 9: NMR titration of SL1-dI into Hexim1 BR-L-AR. A** $^{15}$N-$^{1}$H HSQC spectral overlay with at 600MHz of BR-L-AR (red) with addition of SL1-dI at 1:0.5 (orange), 1:0.8 (cyan) and 1:2 (black) molar ratios. **B** Chemical shift perturbations (CSP) of BR-L-AR upon binding SL1-dI. **C** Intensity ratio of BR-L-AR bound to SL1-dI versus free BR-L-AR.

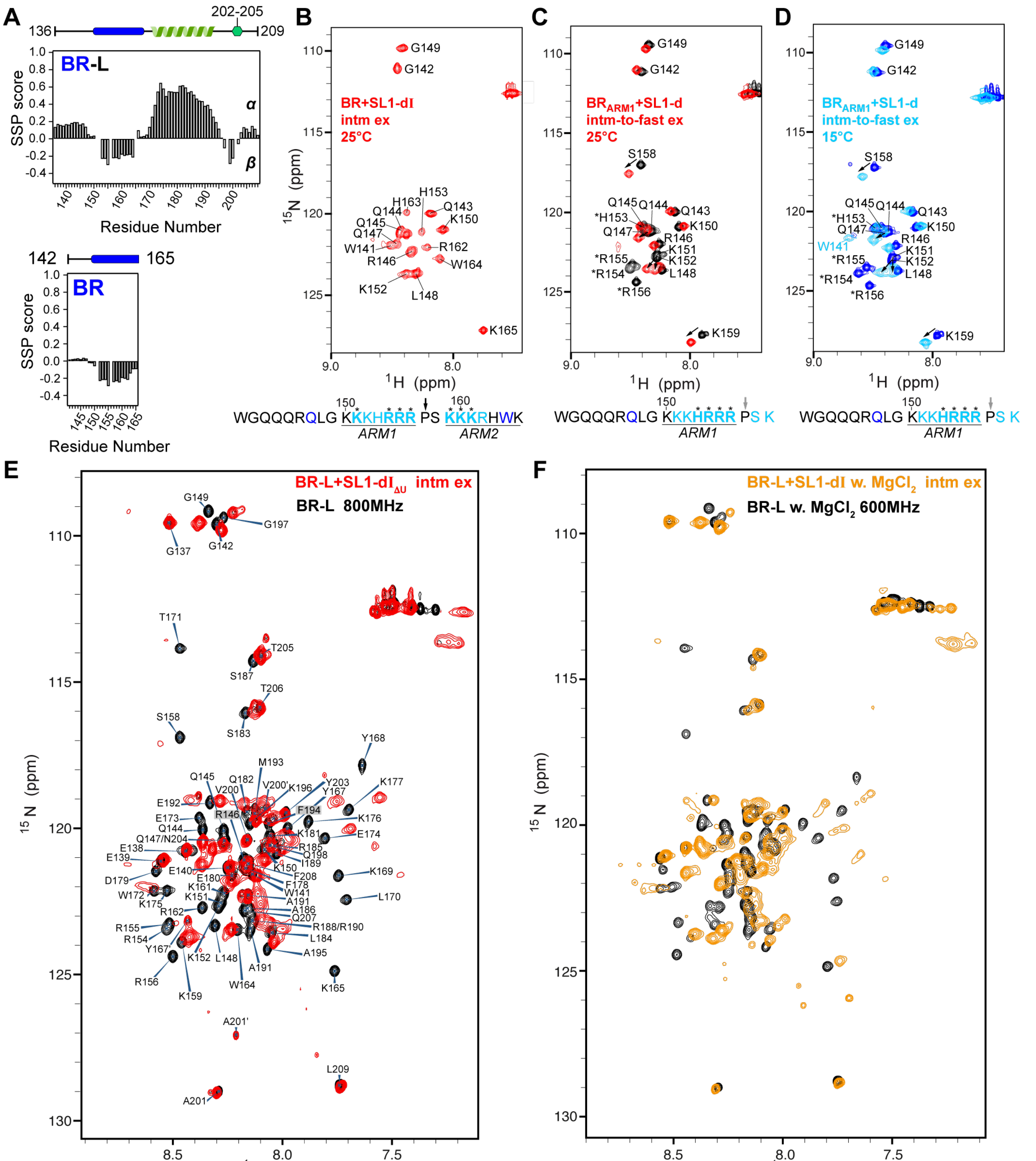


**Supplementary Fig. 10: Spectral behavior of Hexim1 BR-L, BR and $BR_{ARM1}$ upon binding 7SK RNA. A** Secondary structural propensity (SSP) scores of Hexim1 BR-L and BR. **B–D** $^{15}N$-$^{1}H$ HSQC spectra with assignments of Hexim1 BR bound to SL1-dI at 25°C (B), $BR_{ARM1}$ in the absence (black) and bound to SL1-d at 25°C (red) (C) or 15°C (black versus blue) (D). The amino acid sequence of BR or $BR_{ARM1}$ is shown below each spectrum, where cyan residues indicate those with above [mean+σ] CSPs or broadened beyond detection (bold cyan with *) upon Site1 binding and blue residues indicate those with above [mean] CSPs. **E** $^{15}N$-$^{1}H$ HSQC spectral overlay with assignments of Hexim1 BR-L free (black) and bound to SL1-$dI_{\Delta U}$ RNA (red, 1:1 molar ratio). **F** $^{15}N$-$^{1}H$ HSQC spectral overlay with assignments of Hexim1 BR-L free (black) and SL1-dI bound (orange, 1:1.25 molar ratio) in the presence of 3mM $MgCl_2$.

**Notes about line broadening due to RNA binding**

In this figure, together with main Fig. 2D,E and Supplementary Fig. 5, the severe line broadening of BR residues upon RNA binding persists in the range of 0–300 mM KCl and different temperatures (Supplementary Figs. 5, 6 and 10). Similar to BR-L-AR, the $BR_{ARM1}$, BR and BR-L constructs exhibited intermediate exchange with significant broadening of BR residues upon binding RNA (Supplementary Fig. 10). These data indicate chemical exchange between two or more RNA-bound protein conformations. Possibilities include interaction with ARM1 or ARM2 or ARM1-ARM2, likely due to both RNA dynamics and IDR ensemble. Consistent with this, we observed two or three distinct sets of resonances in the $^{15}N$-$^{1}H$ HSQC spectrum of the SL1-dI–BR-L with 3mM $Mg^{2+}$ (Supplementary Fig. 10F).

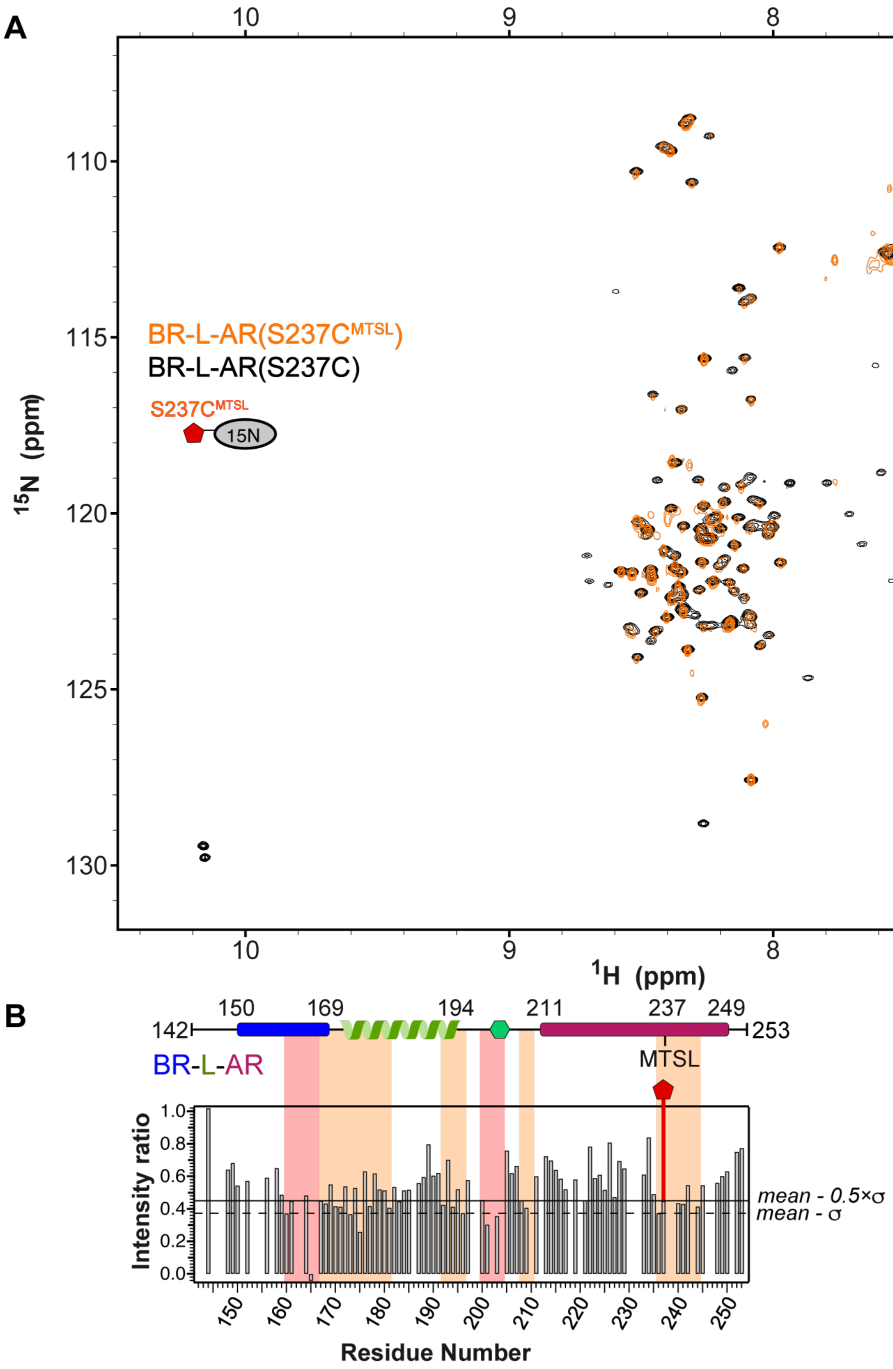


**Supplementary Fig. 11: $^{15}$N BR-L-AR(S237C$^{MTSL}$) broadens more severely than S226C$^{MTSL}$ throughout the protein sequence**. **A** $^{15}$N-$^{1}$H HSQC spectral overlay of Hexim1 BR-L-AR(S237C) variant in the absence (black) and presence (orange) of MTSL tag modification. **B** PRE intensity ratios ($I_{PRE}/I_{noPRE}$) of resolved resonances plotted against residue number. Solid line indicates value of [mean-0.5×σ] and dashed line indicates value of [mean-σ]. Stretches of residues with intensity ratios below the dashed line or below solid lines are highlighted with red and orange shaded boxes, respectively. Schematic of BR-L-AR construct with the S237C$^{MTSL}$ tag location highlighted is shown at top.

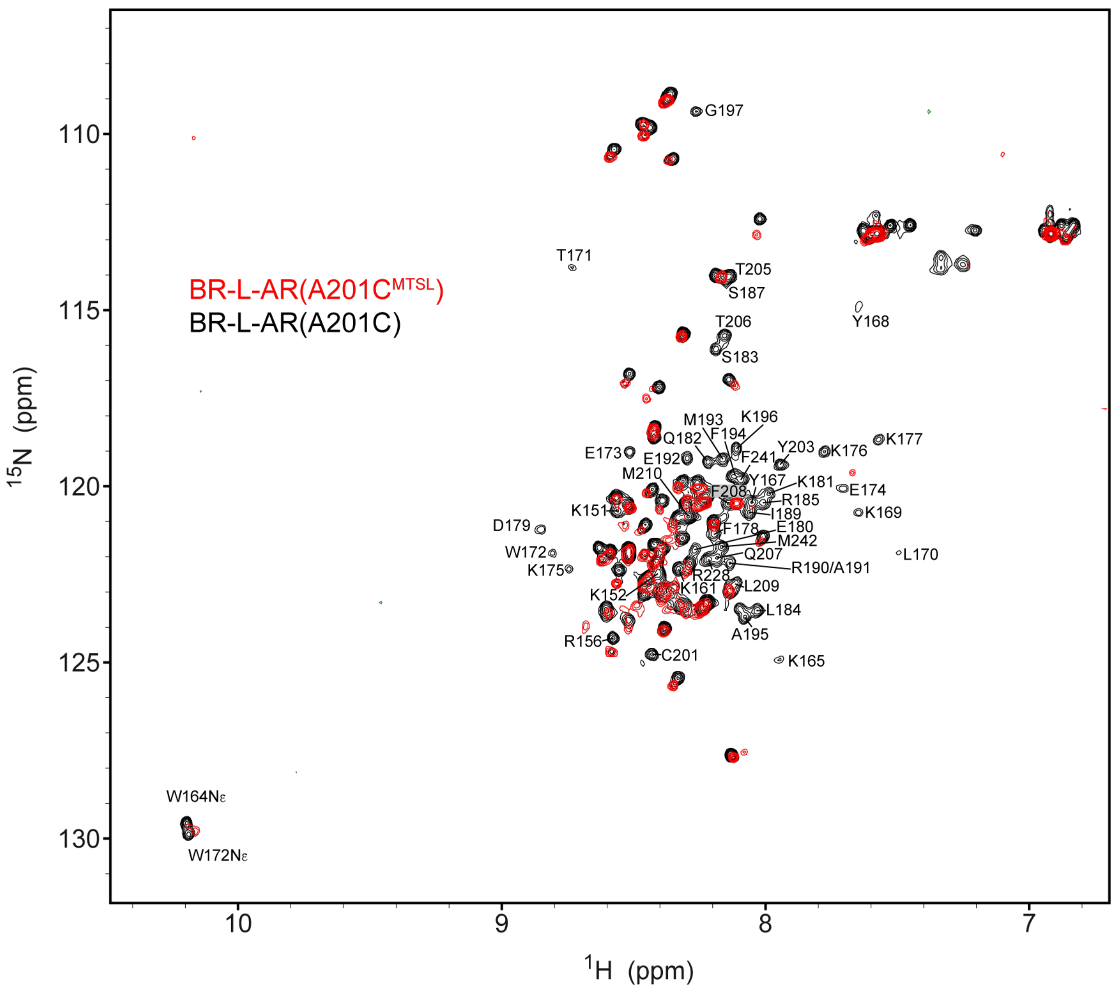


**Supplementary Fig. 12: $^{15}N$-$^{1}H$ HSQC spectral overlay of $^{15}N$ BR-L-AR(A210C)** variant in the absence (black) and presence (red) of MTSL tag modification. Residues that are significantly broadened are labeled on the spectrum.

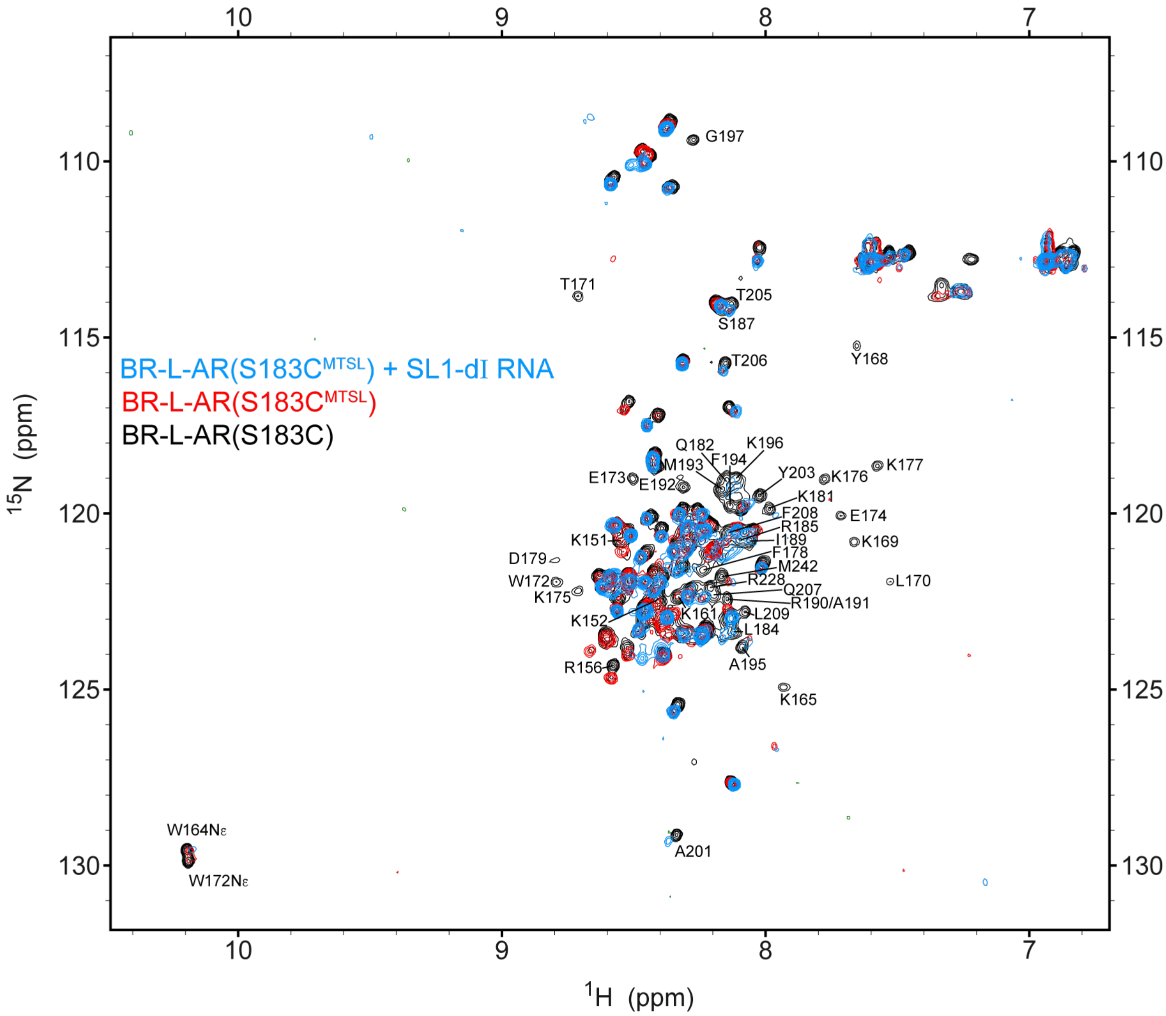


**Supplementary Fig. 13: $^{15}N$-$^{1}H$ HSQC spectral overlay of $^{15}N$ BR-L-AR(S183C)** variant in the absence (black) and presence (red) of MTSL tag modification, and with MTSL modification bound to SL1-dI RNA (cyan). Residues that are significantly broadened are labeled on the spectrum.

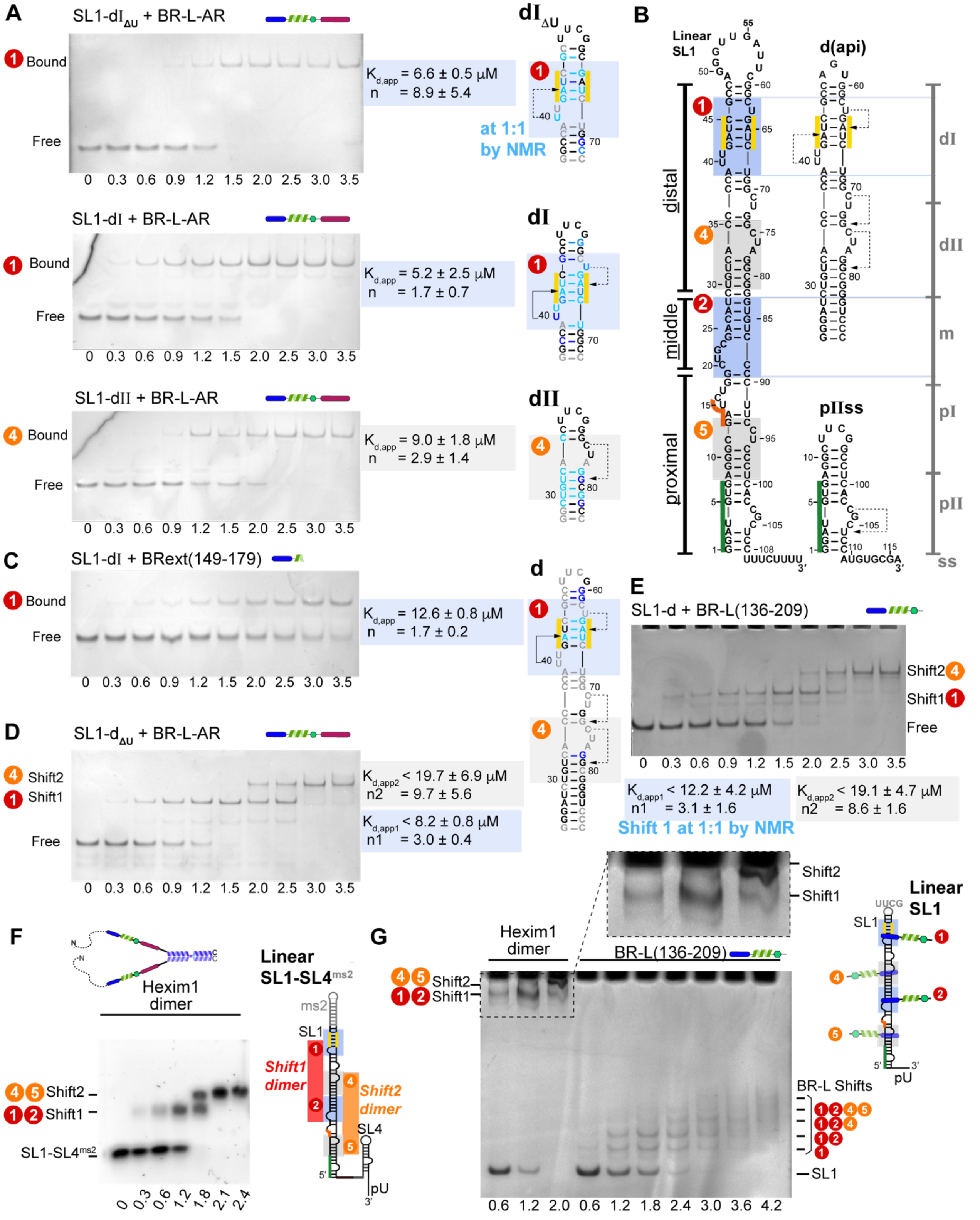

**Supplementary Fig. 14: Binding studies of SL1 RNA with Hexim1 by EMSA. A** Electrophoretic mobility shift assay (EMSA) of Site1 and Site4 constructs binding to Hexim1 BR-L-AR. Protein-to-RNA molar ratios are indicated at the bottom of each gel image. Sites that correspond to each shift event are indicated to the left of each gel. Apparent $K_{D,app}$ and n values are shaded according to the binding sites mapped by NMR, shown to the right. **B** Sequences and secondary structures of additional RNA constructs used in this study. **C** BRext(149–179) exhibits weaker binding than BR-L-AR to SL1-dI by EMSA. **D** Longer construct SL1-$d_{\Delta U}$ has two binding events by EMSA. For these two sequential shifts, the first shift is assigned to Site1 based on NMR CSP of 1-to-1 molar ratio binding experiment, where Site1 exhibit CSP instead of Site4. The saturating ratios of these two shifts are also consistent with the EMSA saturating ratios of individual RNA sites, e.g. both SL1-dI$_{\Delta U}$ shift and the Shift1 of SL1-d$_{\Delta U}$ saturates at 1.5:1, whereas Shift2 of SL1-d$_{\Delta U}$ saturates at 3:1 (2:1 after subtracting the first protein molecule), corresponding to the 2.5:1 saturating ratio of SL1-dII. **E** Longer construct SL1-d has two binding events with BR-L by EMSA. **F** EMSA using an agarose gel for linear SL1-SL4$^{ms2}$ RNA binding to Hexim1 homodimer. Molar ratios are dimer:RNA for full-length Hexim1. This RNA construct is a functionally minimum unit for assembling 7SK RNP with MePCE, Larp7, Hexim1 and P-TEFb in cells[3]. **G** EMSA for full-length linear 7SK SL1 binding to Hexim1 homodimer or BR-L monomer. A zoom-in view of the Hexim1 dimer shifts is shown at the top. Molar ratios are dimer:RNA for full-length Hexim1, and monomer:RNA for BR-L. The multiple shifts of BR-L illustrate that linear SL1 can simultaneously bind four or more BR-L monomers at 4.2:1 protein-to-RNA molar ratio.

**Notes for EMSA binding curve fittings**

The binding behavior on EMSAs all appear to be highly cooperative, likely due to influence of electric field on highly negatively charged RNA and highly positively charged protein molecules. The concentrations used in PAGE-based EMSAs are 9 μM for RNA and 0–32 μM for proteins, in order to match conditions of NMR experiments to determine binding stoichiometry. These concentrations are not in the quantitative range of concentrations to determine $K_D$ values (concentrations of RNA should be below $K_D$ value, which is in the range of ~83–336 nM according to our ITC results). Instead, we acquired apparent $K_D$ values, which are ligand concentrations producing half occupation, or sometimes referred to as microscopic dissociation constants, to strictly compare across EMSA results, which cannot be used to compare with ITC results.

For fitting individual binding sites (SL1-dI$_{\Delta U}$, SL1-dI and SL1-dII), the apparent $K_D$ values were determined by fitting fraction bound using both free RNA bands and protein-bound bands individually to the following equation:

$$F = \frac{A\left([Pt] - \frac{[Rt]+[Pt]+K_{D,app} - \sqrt{\left([Rt]+[Pt]+K_{D,app}\right)^2 - 4[Rt][Pt]}}{2}\right)^n}{K_{D,app}^n + \left([Pt] - \frac{[Rt]+[Pt]+K_{D,app} - \sqrt{\left([Rt]+[Pt]+K_{D,app}\right)^2 - 4[Rt][Pt]}}{2}\right)^n}$$

where *[Rt], [Pt],* $K_{D,app}$*, n, F* and *A* are the total RNA concentration, protein concentration, apparent dissociation constant, the Hill coefficient, fraction bound and a correction factor for maximum fraction bound, respectively. This equation accounts for the bound portion of protein and subtracts them from the total protein concentrations at each titration point, in order to yield free protein concentrations.

For fitting two sequential binding sites for SL1-d$_{\Delta U}$ and SL1-d, an approximation of using total protein concentration as free protein concentration was used, in order to still include Hill coefficients for each site using the following equation to yield converging fitting results:

$$F0 = A0\left(1 - \frac{K_{D,app1}^{n1}K_{D,app2}^{n2}}{K_{D,app1}^{n1}K_{D,app2}^{n2} + K_{D,app2}^{n2}[Pt]^{n1} + [Pt]^{n1+n2}}\right)$$

$$F1 = A1\left(\frac{K_{D,app2}^{n2}[Pt]^{n1}}{K_{D,app1}^{n1}K_{D,app2}^{n2} + K_{D,app2}^{n2}[Pt]^{n1} + [Pt]^{n1+n2}}\right)$$

$$F1 = A2\left(\frac{[Pt]^{n1+n2}}{K_{D,app1}^{n1}K_{D,app2}^{n2} + K_{D,app2}^{n2}[Pt]^{n1} + [Pt]^{n1+n2}}\right)$$

where *[Pt]*, $K_{D,app1}$, $K_{D,app2}$, *n1* and *n2* are the total protein concentration, apparent dissociation constants and the Hill coefficients for first site and second site, respectively. *F0, F1* and *F2* are fractions for [1 - free RNA] (for display purpose), one protein molecule bound RNA and two protein molecules bound RNA, respectively. *A0, A1* and *A2* are correction factors for maximum fraction bound. The free bands, Shift1 and Shift2 bands are fit independently, and resulting values averaged. The $K_{D,app1}$ and $K_{D,app2}$ values are over-estimated due to having to use the approximation of total protein concentration as the free protein concentration.

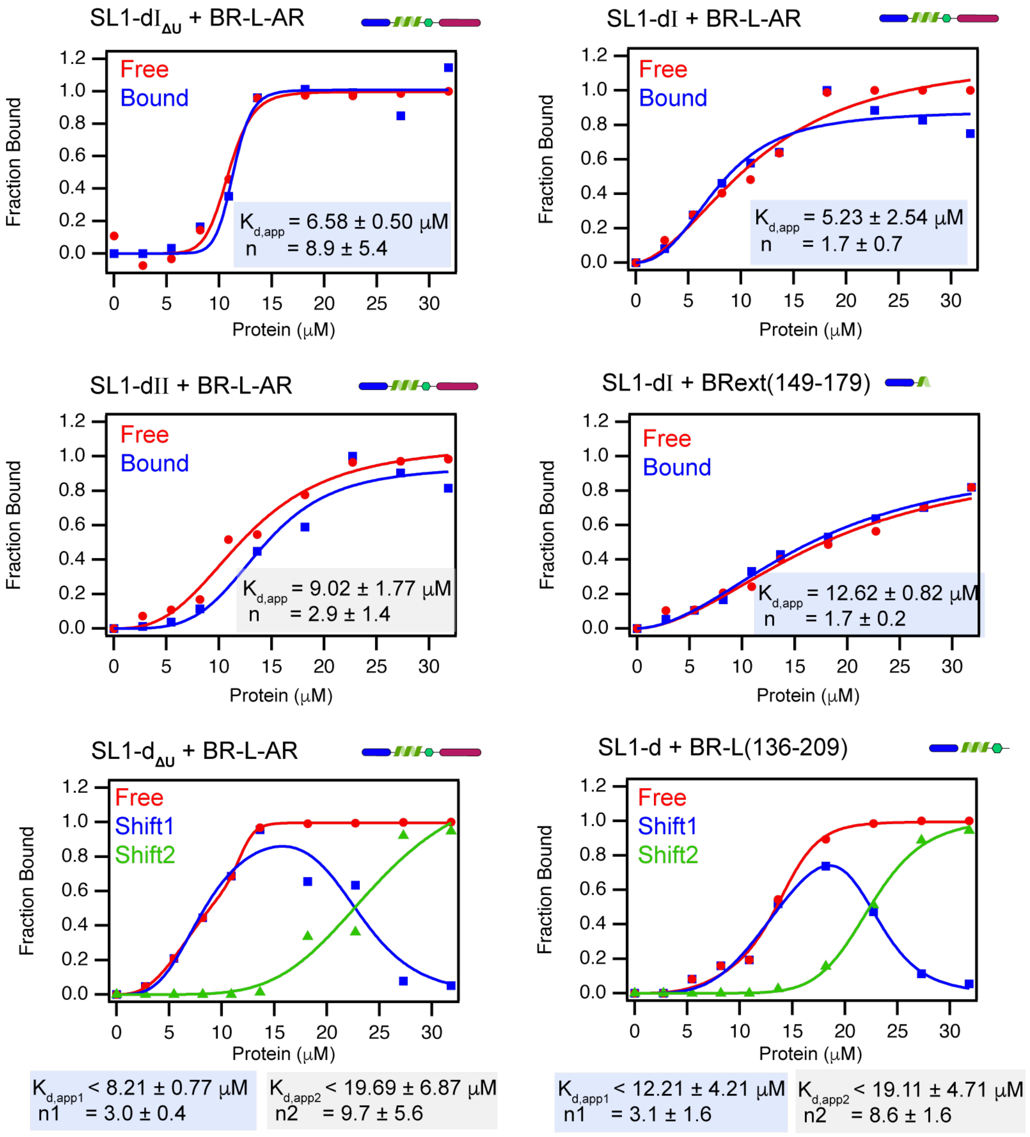


**Supplementary Fig. 15: Fitting results from EMSA studies.** See above notes for fitting details. For fitting two sequential binding sites for SL1-d$_{ΔU}$ and SL1-d (bottom two panels), the $K_{D,app1}$ and $K_{D,app2}$ values are over-estimated due to having to use the approximation of total protein concentration as the free protein concentration, hence the acquired apparent dissociation constant values from fittings are indicated as the upper bounds.

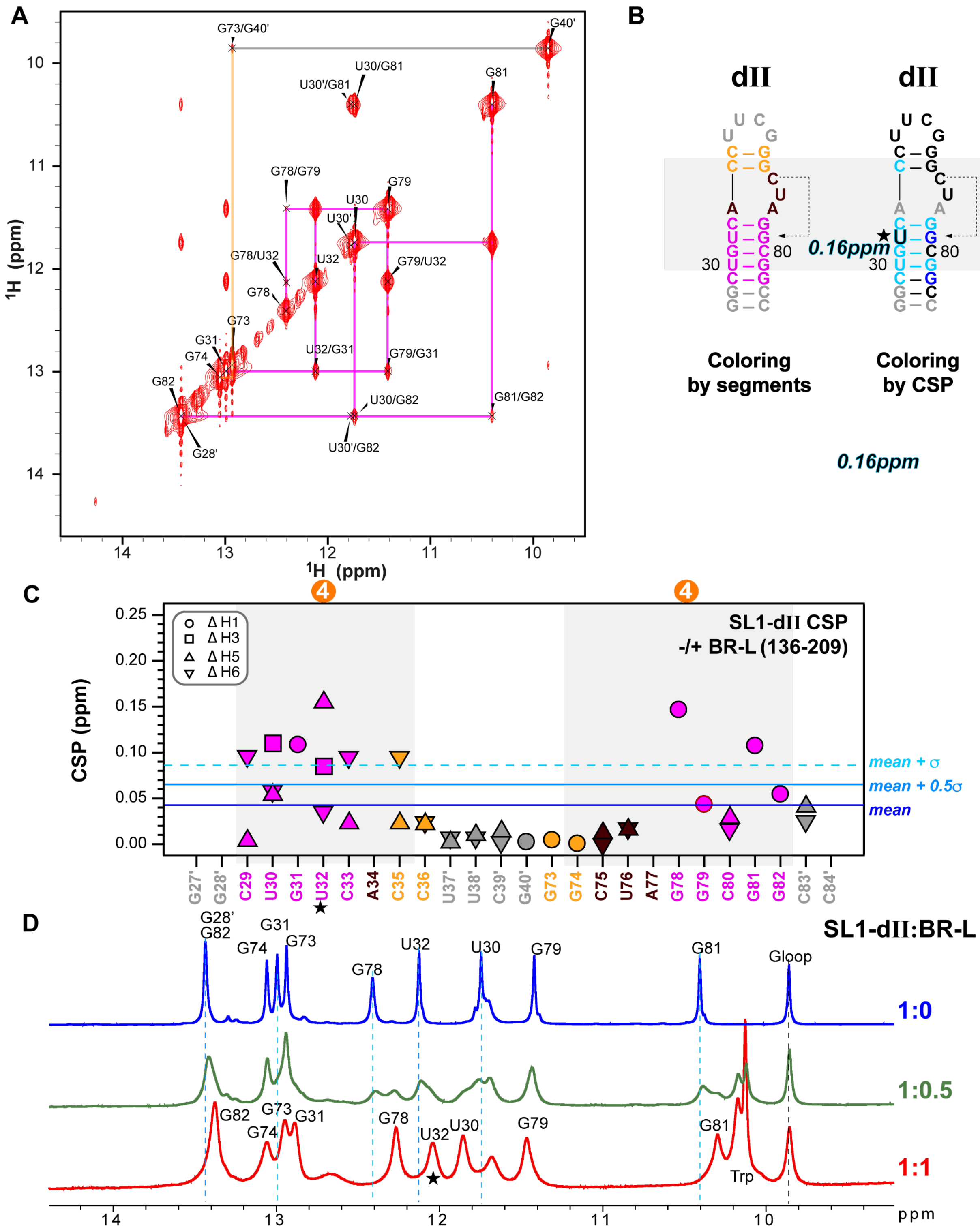


**Supplementary Fig. 16: NMR binding studies of SL1-dII RNA with Hexim1 BR-L. A** 2D imino NOESY assignments of SL1-dII RNA. **B** Secondary structure of SL1-dII RNA colored by segment and by Chemical shift perturbations (CSP) upon binding BR-L, respectively. The nucleotide experiencing the largest shift is highlighted with black/cyan font and a star, with the corresponding CSP value labeled. **C** CSP plot of SL1-dII RNA upon binding BR-L. Severely broadened and highly overlapping resonances were excluded from the free/bound pair for CSP calculations. **D** 1D imino overlay of SL1-dII in the absence and presence of BR-L.

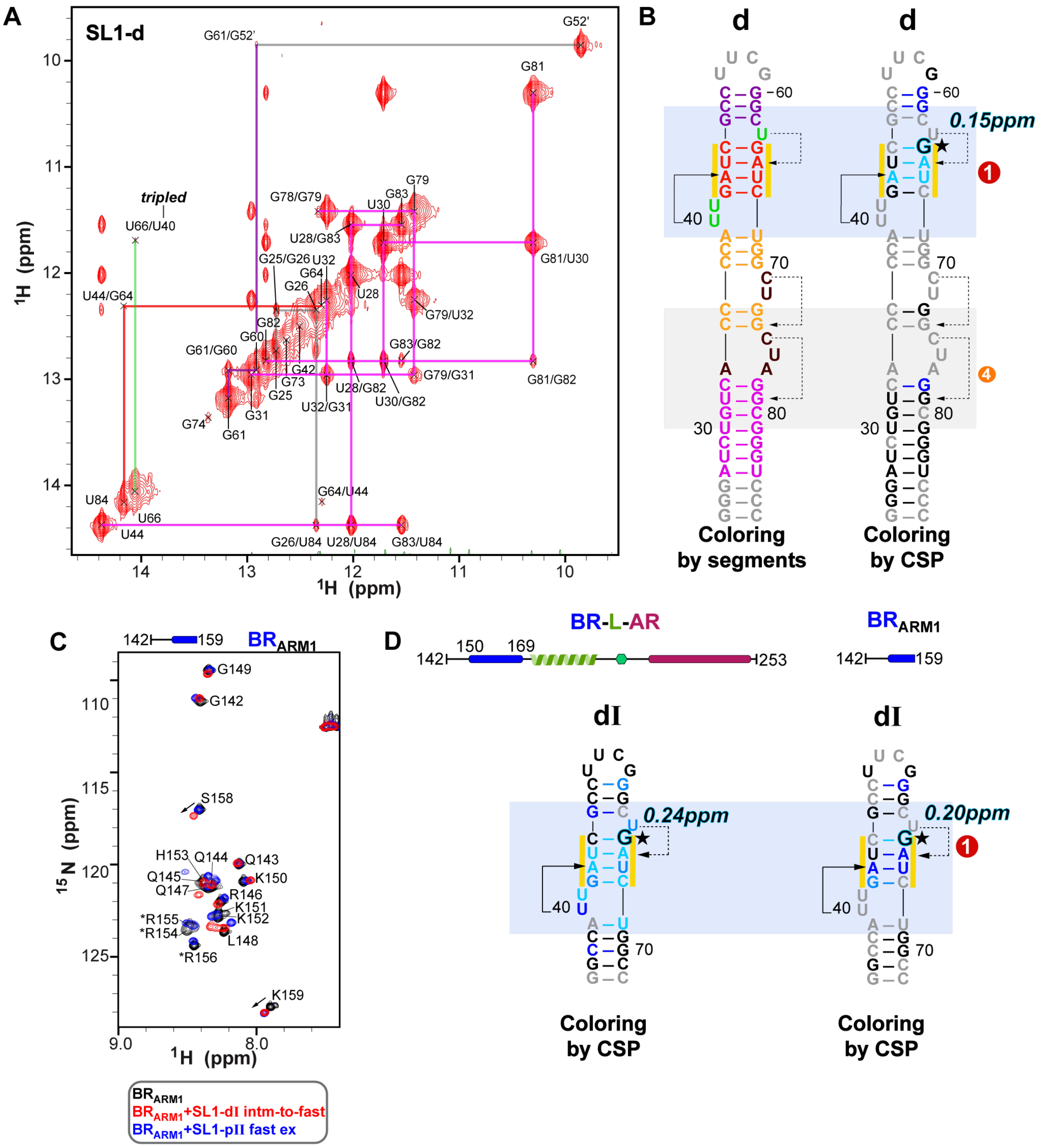

**Supplementary Fig. 17: NMR binding studies of SL1-d RNA with Hexim1 BR$_{ARM1}$. A** 2D imino NOESY assignments of SL1-d RNA. **B** Secondary structure of SL1-d RNA colored by segment and by Chemical shift perturbations (CSP) upon binding BR$_{ARM1}$, respectively. **C** $^{15}$N-$^{1}$H HSQC spectral overlay of Hexim1 BR$_{ARM1}$ (black) and with addition of 1:1 SL1-dI (red; high affinity, intermediate-to-fast exchange) or 1:1 SL1-pII RNA (blue; non-specific site, fast exchange), likely due to a faster off-rate for SL1-pII. **D** CSP of SL1-dI RNA upon binding Hexim1 BR-L-AR versus BR$_{ARM1}$. In (B) and (D), the nucleotide experiencing the largest shift is highlighted with black/cyan font and a star, with their corresponding CSP values labeled. Severely broadened and highly overlapping resonances were excluded from the free/bound pair for CSP calculations.

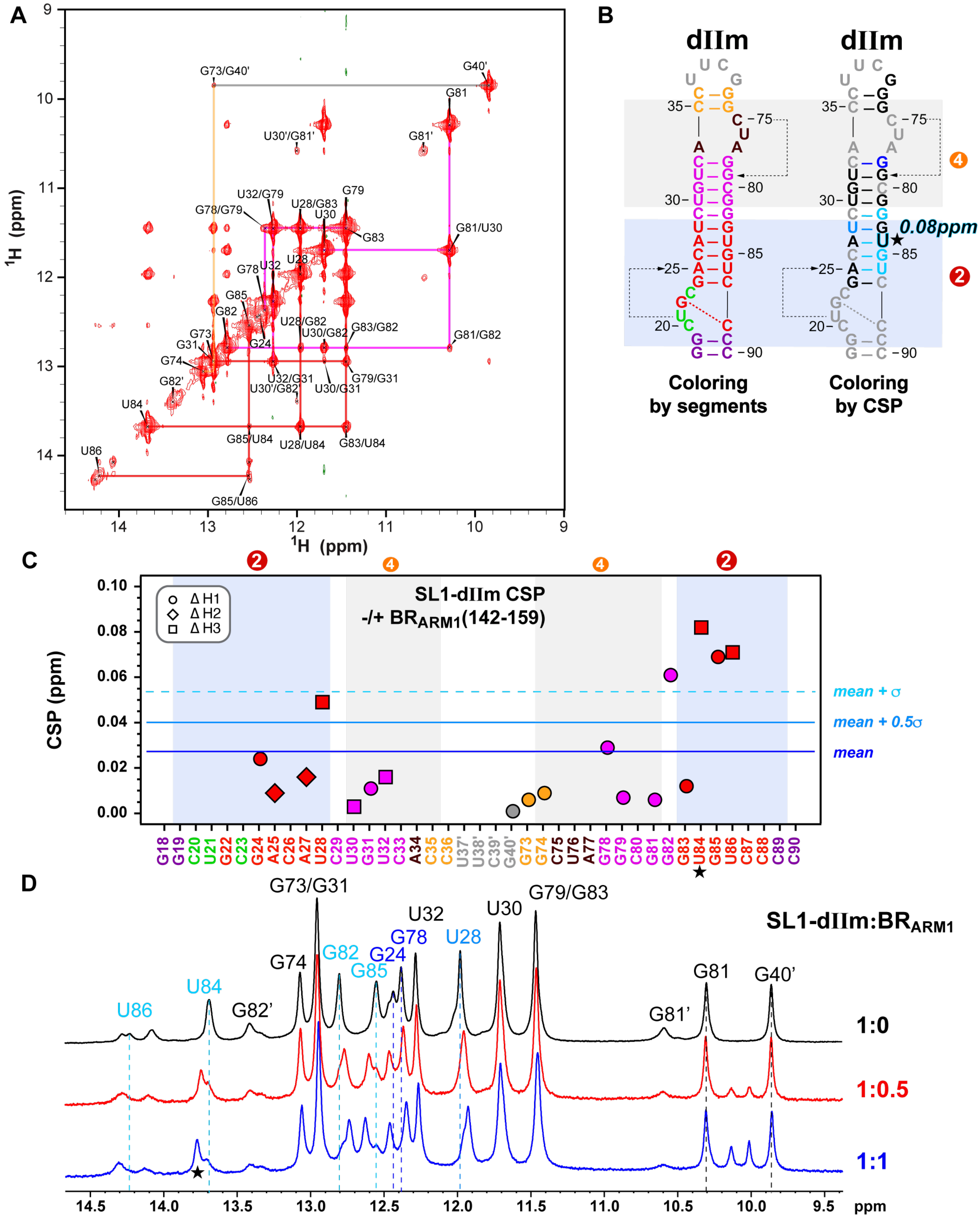

**Supplementary Fig. 18: NMR binding studies of SL1-dIIm RNA with Hexim1 $BR_{ARM1}$. A** 2D imino NOESY assignments of SL1-dIIm RNA. **B** Secondary structure of SL1-dIIm RNA colored by segment and by Chemical shift perturbations (CSP) upon binding $BR_{ARM1}$, respectively. The nucleotide experiencing the largest shift is highlighted with black/cyan font and a star, with the corresponding CSP value labeled. **C** CSP plot of SL1-dIIm RNA upon binding $BR_{ARM1}$. Severely broadened and highly overlapping resonances were excluded from the free/bound pair for CSP calculations. **D** 1D imino overlay of SL1-dIIm in the absence and presence of $BR_{ARM1}$.

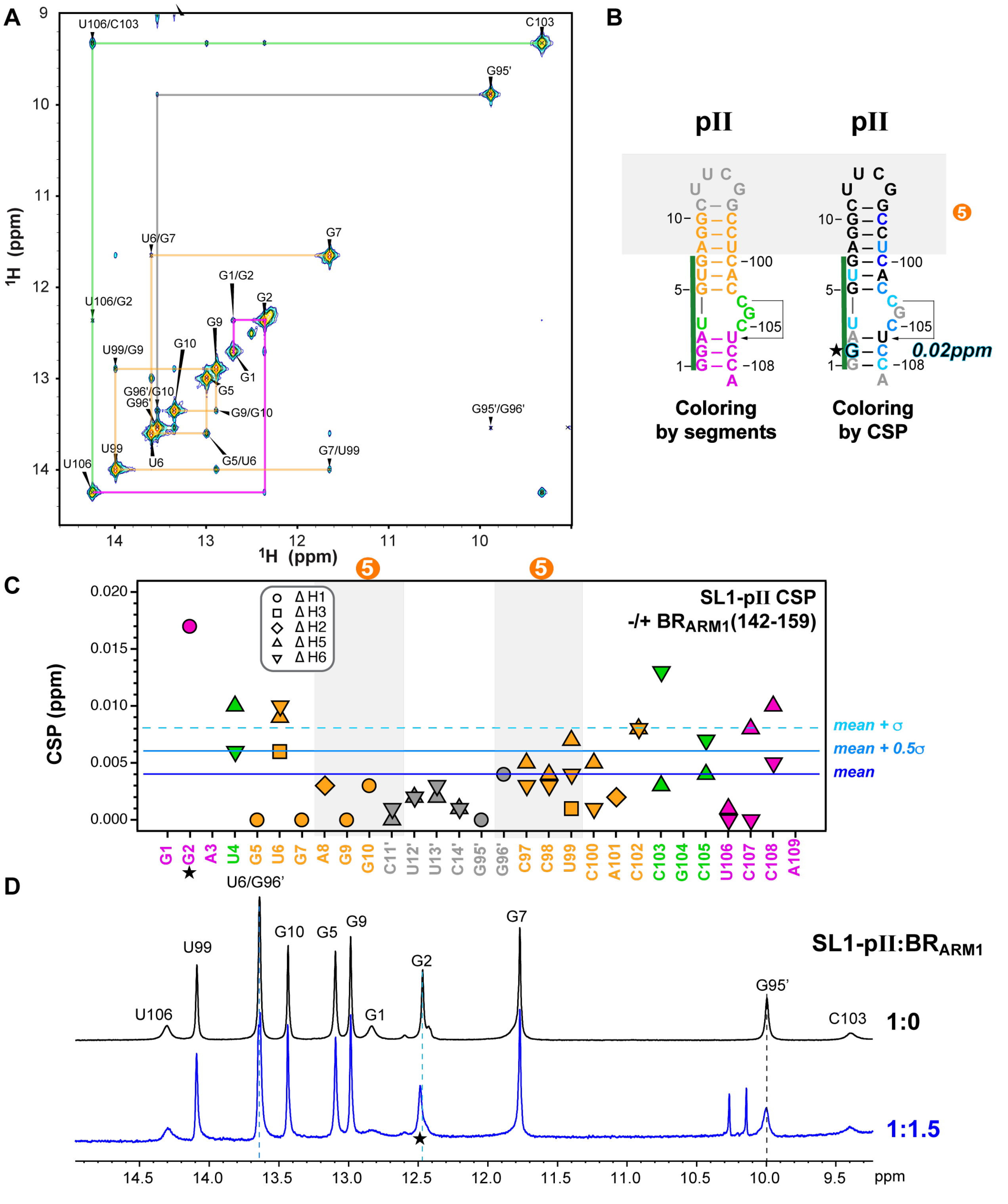


**Supplementary Fig. 19: NMR binding studies of SL1-pII RNA with Hexim1 BR$_{ARM1}$. A** 2D imino NOESY assignments of SL1-pII RNA. **B** Secondary structure of SL1-pII RNA colored by segment and by Chemical shift perturbations (CSP) upon binding BR$_{ARM1}$, respectively. The nucleotide experiencing the largest shift is highlighted with black/cyan font and a star, with the corresponding CSP value labeled. Note that SL1-pII RNA construct only contains the stem of Site 5, lacking the $C_{94}U_{95}$ bulge that forms the base triple in the complete Site 5 binding region. **C** CSP plot of SL1-pII RNA upon binding BR$_{ARM1}$. Severely broadened and highly overlapping resonances were excluded from the free/bound pair for CSP calculations. **D** 1D imino overlay of SL1-pII in the absence and presence of BR$_{ARM1}$.

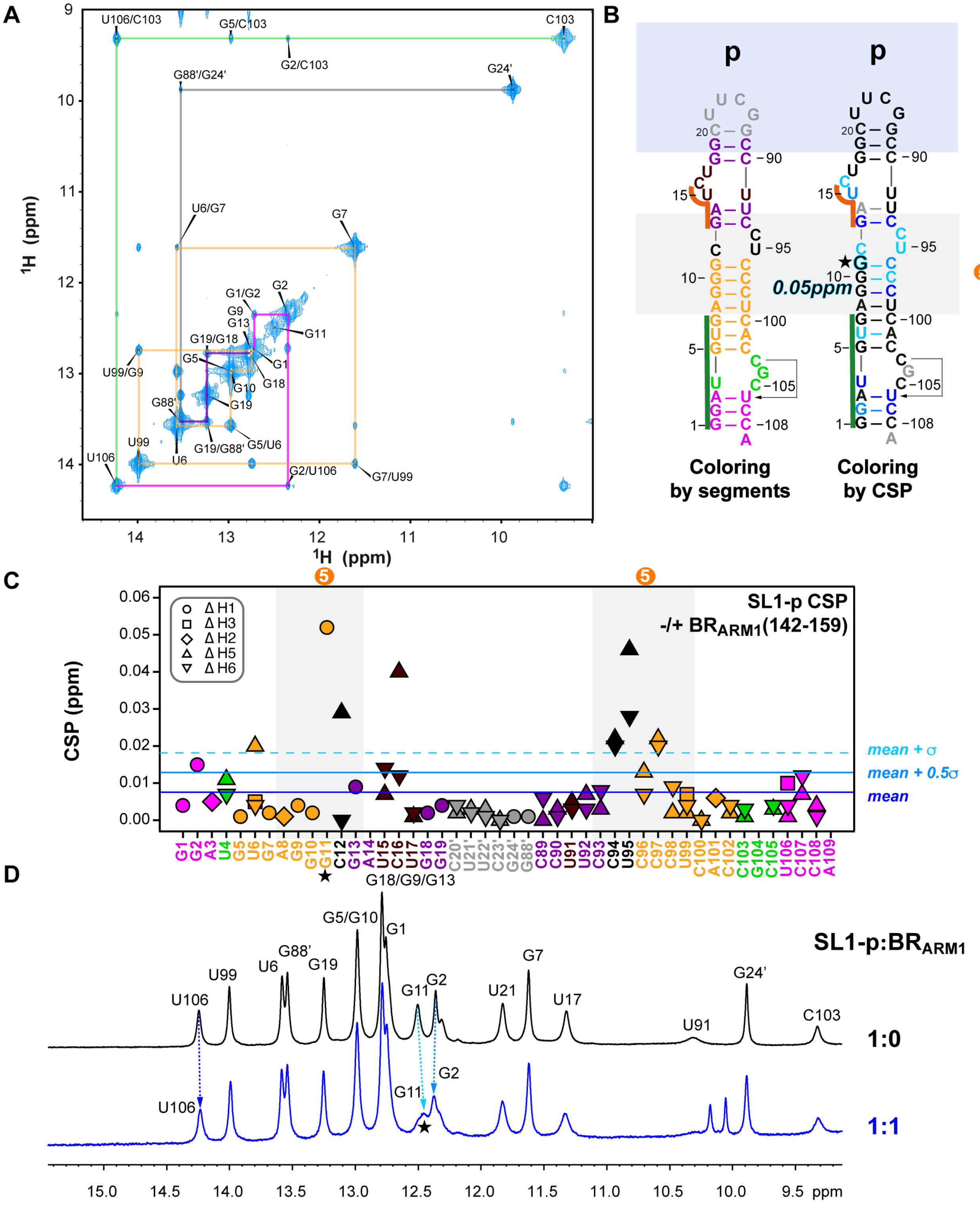

**Supplementary Fig. 20: NMR binding studies of SL1-p RNA with Hexim1 BR$_{ARM1}$. A** 2D imino NOESY assignments of SL1-p RNA. **B** Secondary structure of SL1-p RNA colored by segment and by Chemical shift perturbations (CSP) upon binding BR$_{ARM1}$, respectively. The nucleotide experiencing the largest shift is highlighted with black/cyan font and a star, with the corresponding CSP value labeled. **C** CSP plot of SL1-p RNA upon binding BR$_{ARM1}$. Severely broadened and highly overlapping resonances were excluded from the free/bound pair for CSP calculations. **D** 1D imino overlay of SL1-p in the absence and presence of BR$_{ARM1}$.

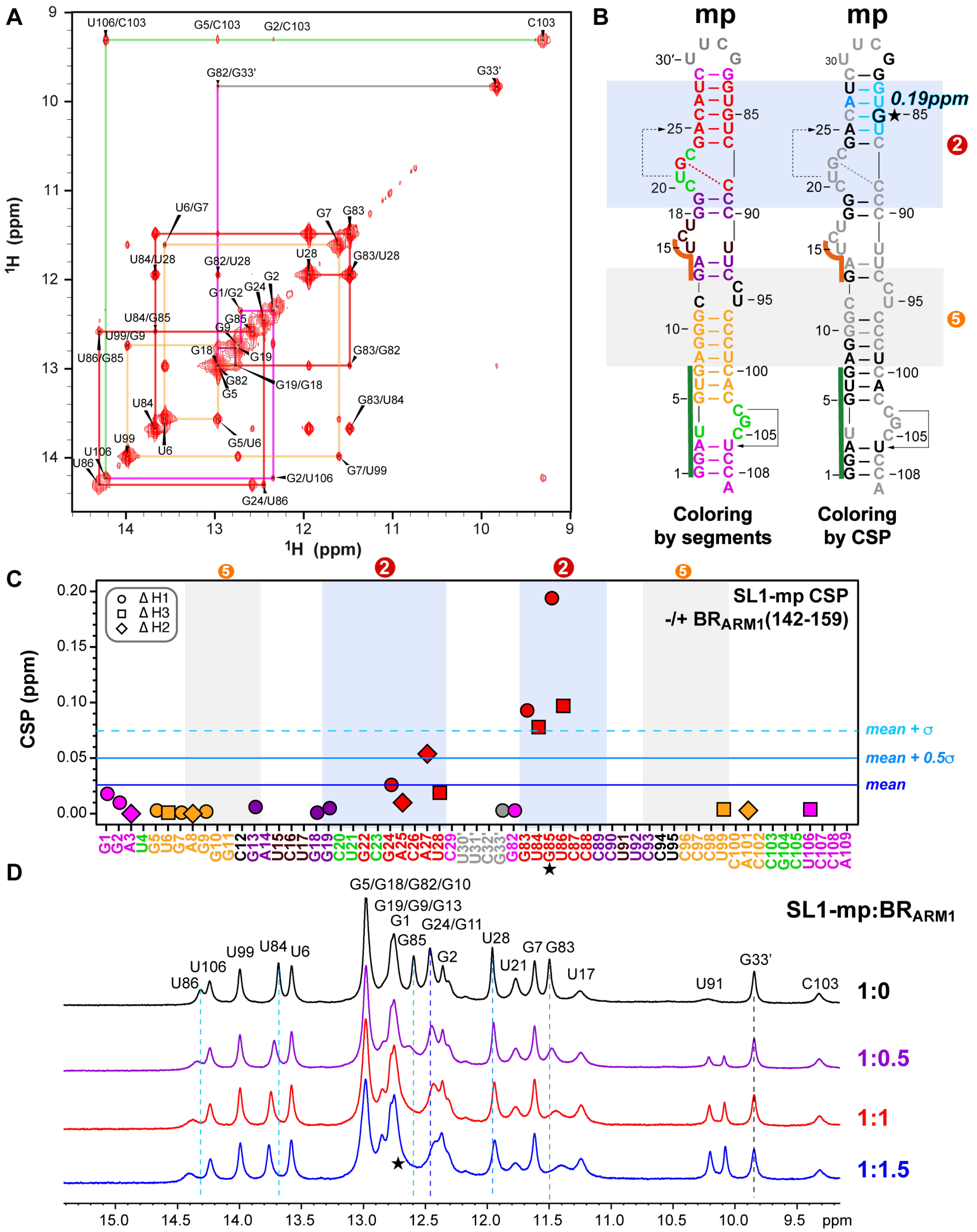

**Supplementary Fig. 21: NMR binding studies of SL1-mp RNA with Hexim1 BR$_{ARM1}$. A** 2D imino NOESY assignments of SL1-mp RNA. **B** Secondary structure of SL1-mp RNA colored by segment and by Chemical shift perturbations (CSP) upon binding BR$_{ARM1}$, respectively. The nucleotide experiencing the largest shift is highlighted with black/cyan font and a star, with the corresponding CSP value labeled. **C** CSP plot of SL1-mp RNA upon binding BR$_{ARM1}$. **D** 1D imino overlay of SL1-mp in the absence and presence of BR$_{ARM1}$. Note that the RNA was already saturated at 1:1, as no further change of RNA peaks was observed between 1:1 and 1:1.5 spectra.

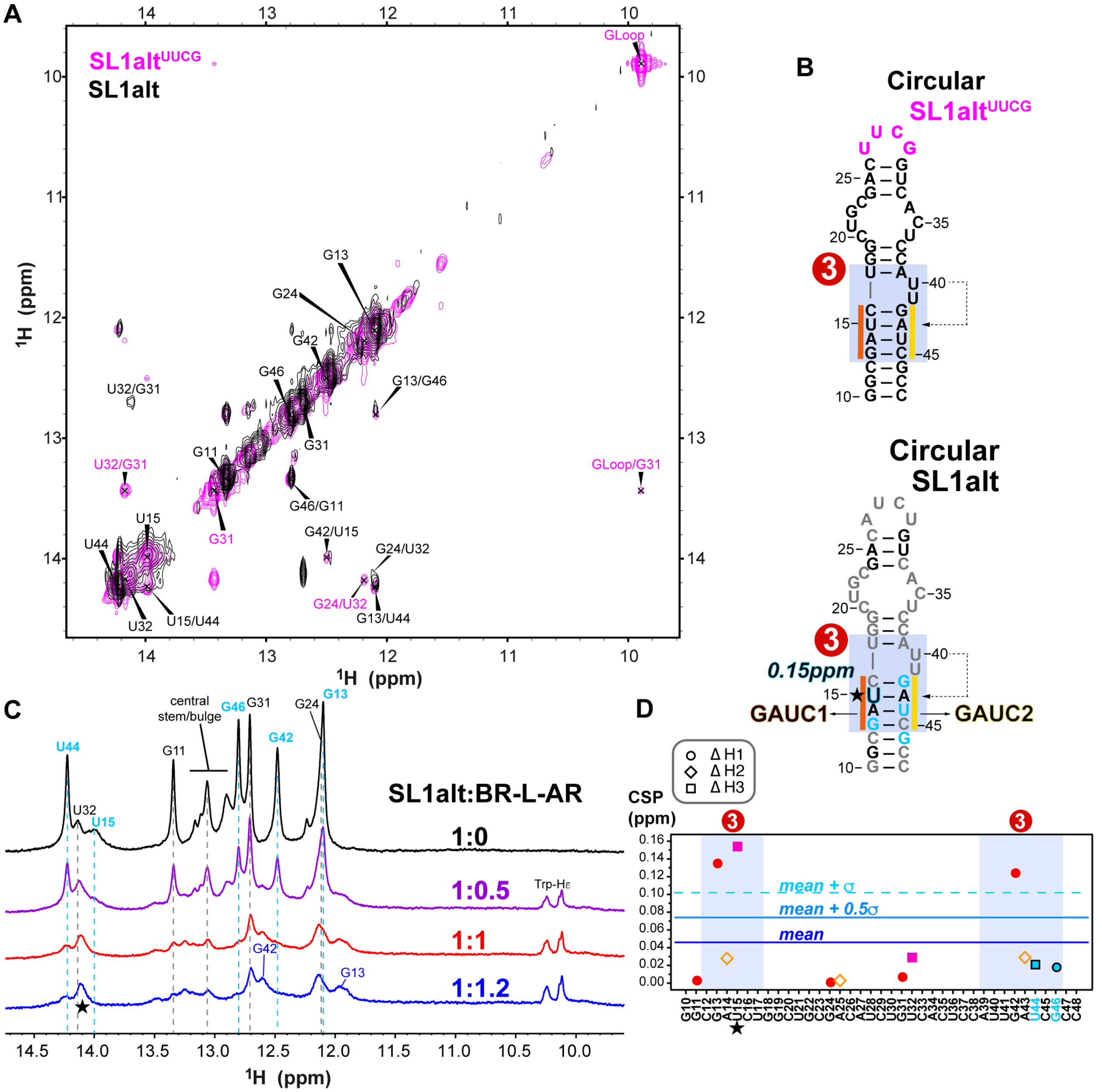

**Supplementary Fig. 22: NMR assignments and binding study of circular 7SK SL1alt RNA at 800MHz. A** Imino region of 2D $^{1}$H-$^{1}$H NOESY spectra with SL1alt (UUCG-tetraloop, magenta) and SL1alt-wt (black) assignments. Assignments for both RNA constructs are shown as peak labels, whereas a couple of SL1alt specific assignments are magenta labels. **B** Secondary structure of SL1alt$^{UUCG}$, which is a UUCG-tetraloop version of SL1alt. **C** Proton 1D imino region of SL1alt RNA in the absence (black) and presence of Hexim1 BR-L-AR (purple, red and blue for 1:0.5, 1:1 and 1:1.2 ratios, respectively). Resonances most affected by protein binding are highlighted with cyan peak labels. **D** CSP plot of SL1alt RNA upon binding Hexim1 BR-L-AR. U44 and G46 are broadened beyond detection, but their chemical shift values of residual RNA-free peak are kept for the purpose of calculating CSP mean and standard deviations. The nucleotide experiencing the largest shift is highlighted with black/cyan font and a star, with the corresponding CSP value labeled.

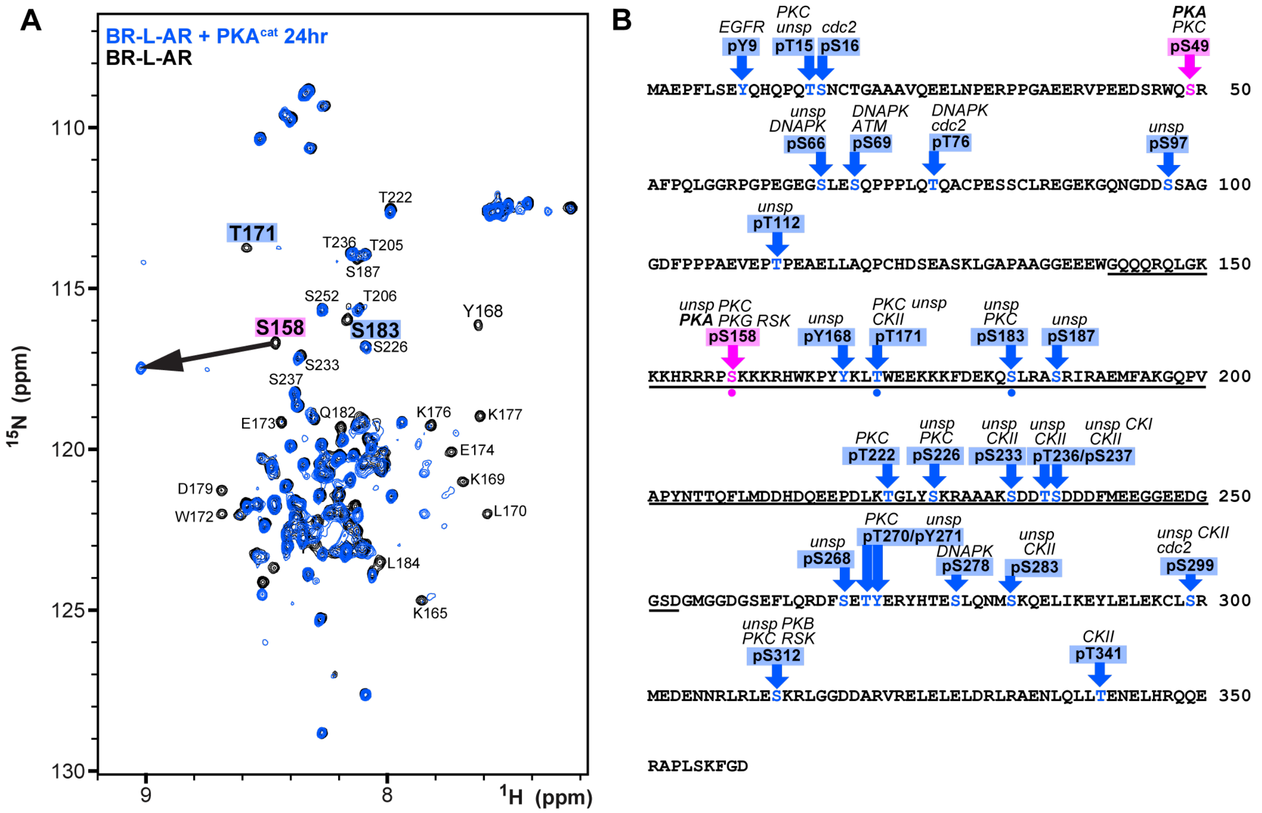


**Supplementary Fig. 23: Longer incubation time reveals secondary phosphorylation sites in αL helix. A** $^{15}$N-$^{1}$H HSQC spectral overlay of BR-L-AR without (black) and with PKA$^{cat}$ incubation at 37°C for 24hr. **B** Predicted phosphorylation sites of Hexim1 by NetPhos-3.1b server[4]. BR-L-AR sequence is underlined, and the three observed sites in our *in vitro* phosphorylation experiment are indicated by dots. Another PKA phosphorylation site in the N-terminus has been predicted (pS49, pink arrow).

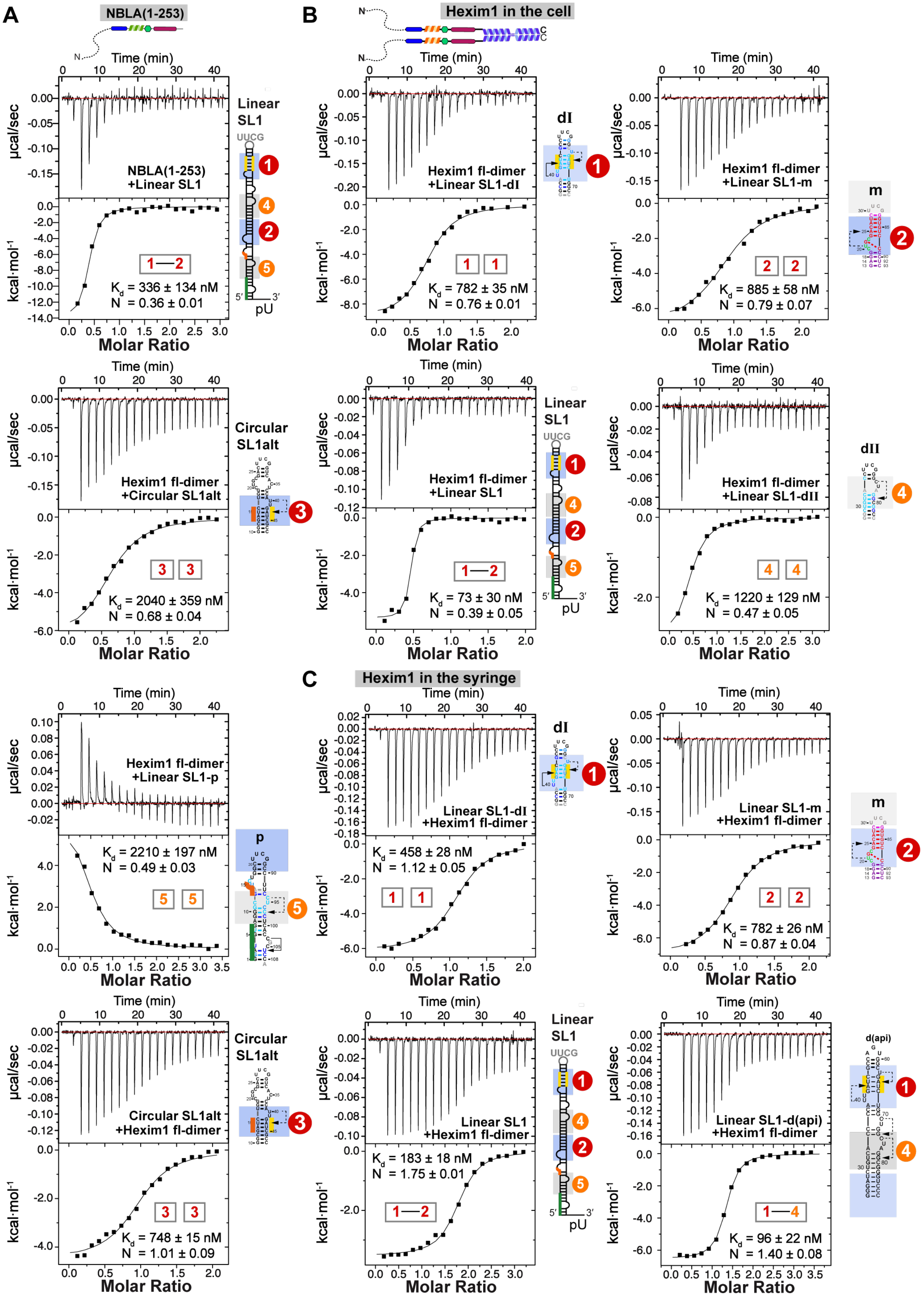


**Supplementary Fig. 24: ITC measurements of NBLA and Hexim1 dimer binding to 7SK RNA constructs. A** Representative ITC measurement of NBLA binding to full-length linear SL1. **B**

Representative ITC measurements of Hexim1 dimer binding to SL1-dI, SL1-m, circular SL1alt (UUCG tetraloop), full-length linear SL1, SL1-dII, and SL1-p. **C** Representative ITC measurements of Hexim1 dimer binding to various RNA constructs using reversed titration order to (B), where Hexim1 dimer protein is in the syringe, and RNA samples are in the sample cell. Secondary structures of RNA constructs are shown to the right of each panel. ITC measurements for each sample were acquired with a minimum of 3 independent titration experiments.

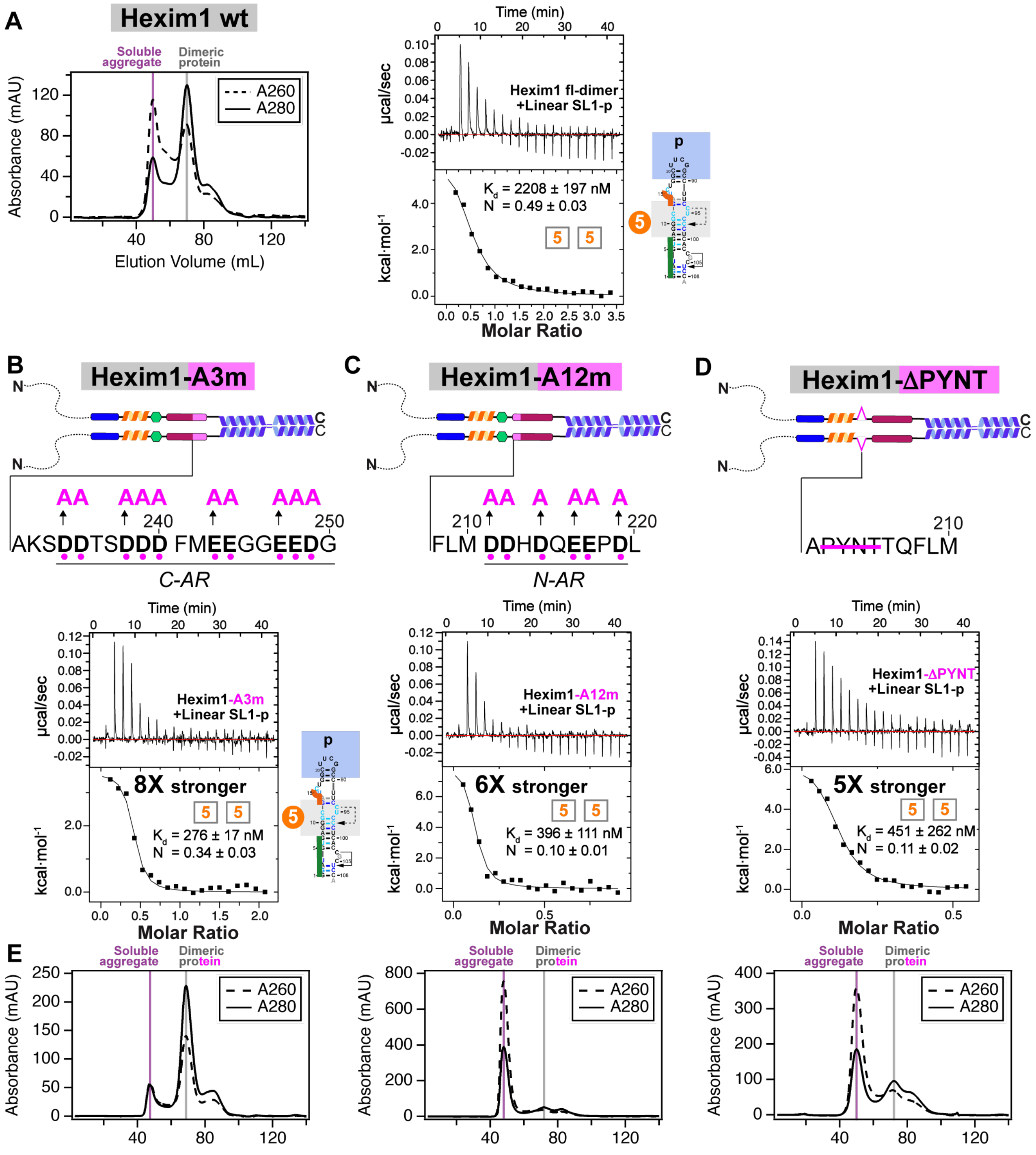


**Supplementary Fig. 25: Hexim1 dimer variants that disrupt autoinhibition became non-specific RNA binder. A** SEC chromatogram and ITC results of wild-type Hexim1 for comparison to variants. **B** Representative ITC measurement of Hexim1-A3m binding to SL1-pII. **C** Representative ITC measurement of Hexim1-A12m binding to SL1-p. **D** Representative ITC measurement of Hexim1-ΔPYNT binding to SL1-p. The affinity fold changes compared to wild-type Hexim1 dimeric protein are indicated next to each ITC results in (B-D). **E** SEC chromatograms of Hexim1 dimer variants during purification. Dimeric protein peak (grey line) fractions were conservatively pooled for ITC, excluding fractions from the soluble aggregate peak (purple line) fractions. Note that A12m and ΔPYNT co-purify as nucleic-acid containing soluble aggregate (higher A260 than A280) much more significantly than wild-type Hexim1 or A3m mutant, indicating stronger non-specific binding to bacterial nucleic acids during purification and/or potential higher propensity to self-aggregate.

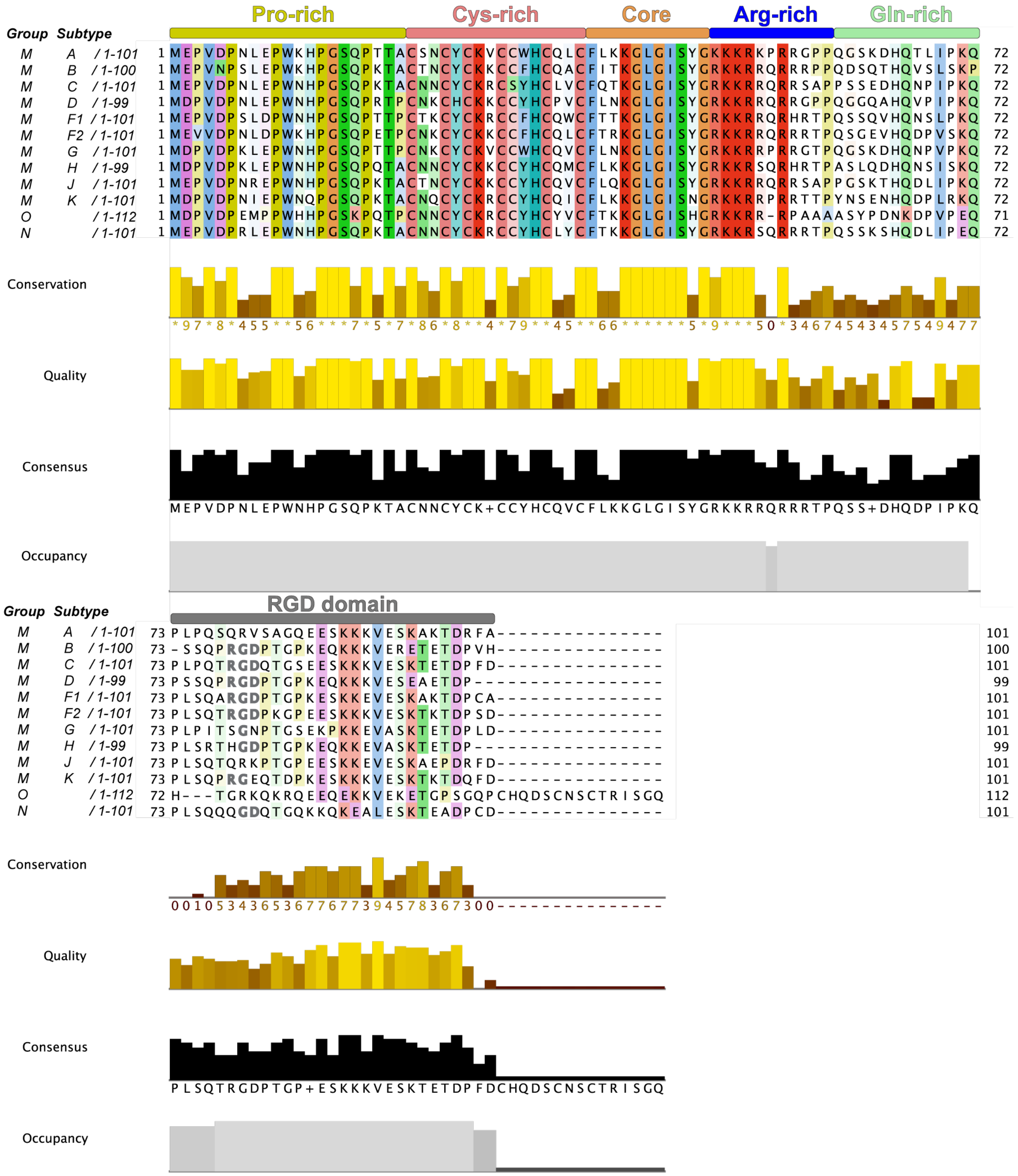


**Supplementary Fig. 26: Sequence alignment of HIV-1 Tat protein.** The viral Groups and Subtypes are indicated to the left of each representative sequence, whose accession IDs are, P24738, Q73370, O12161, P04611, Q9QSR0, P0C1K0, O41801, O70899, Q9WC57, P0C1K4, P0C1K2, O91083, respectively. Only long isoforms with both exons are shown for this display.

**Supplementary discussion regarding N-terminus of Hexim and Tat**

For human Hexim1 sequence, its extreme N-terminus contains a slightly acidic and hydrophobic stretch (residues 1-26, net -4 charge, or -8 charge for residues 1-66), but the acidic residues are not clustered, rather this segment appears more amphipathic than acidic (Supplementary Fig. 1). In comparison, central region AR (-13 total net charges) contains clustered net -6 charges for n-AR, and clustered net -10 charges for c-AR (Supplementary Fig. 2). In a previous biochemical study, a N-terminus deletion construct of Hexim1 exhibited partial premature P-TEFb inhibition activity[5]. Interestingly, this ∆N construct could still respond to 7SK RNA binding, which significantly enhanced its P-TEFb-inhibitory activity, consistent with releasing BR-L-AR mediated autoinhibition. We note that the inclusion of N-terminus increased the solubility of RNP during ITC titrations in our study, pointing to a contribution of the disordered N-terminus to an altered protein biochemical property. It is possible that the extreme N-terminus could provide further stabilization to the BR–PYNT–AR inter-monomer interaction of the central region or contact CC domain.

Similar to Hexim1 N-terminus, HIV-1 Tat has an amphipathic and hydrophobic N-terminus with a non-clustered -2 to -3 net charge (Pro-rich domain, Supplementary Fig. 26). Thus, Tat N-terminus could have a slight autoinhibitory effect to Tat ARM–similar to the N-terminus of Hexim1–pending future investigations.

**Supplementary Table 1: Thermodynamic parameters for binding of Hexim1 to 7SK RNA from ITC with replicate numbers**

| | $K_D$ (nM) | N | ΔH kcal/mol | -TΔS kcal/mol | ΔG kcal/mol | Replicates* |
|---|---|---|---|---|---|---|
| **NBLA monomer** | | | | | | |
| SL1-dI | 83 ± 28 | 0.81 ± 0.10 | -9.2 ± 0.6 | -0.3 ± 0.7 | -9.5 ± 0.2 | 4 |
| SL1-m | 189 ± 25 | 0.62 ± 0.07 | -6.2 ± 0.5 | -2.9 ± 0.4 | -9.1 ± 0.1 | 3 |
| SL1 | 336 ± 134 | 0.36 ± 0.01 | - 14 ± 0 | 5.5 ± 0.1 | -8.8 ± 0.3 | 3 |
| SL1alt** | 261 ± 68 | 0.93 ± 0.06 | -8.9 ± 0.2 | -0.1 ± 0.2 | -9.0 ± 0.2 | 4 |
| ***S158EE*** | | | | | | |
| SL1-dI | 727 ± 63 | 0.74 ± 0.04 | -9.7 ± 0.9 | 1.5 ± 0.9 | -8.3 ± 0.1 | 3 |
| SL1 | 1990 ± 406 | 0.17 ± 0.02 | -27 ± 4 | 19 ± 4 | -7.7 ± 0.1 | 3 |
| **Hexim1 dimer** | | | | | | |
| SL1-dI | 780 ± 25 | 0.76 ± 0.01 | -8.5 ± 0.6 | 0.3 ± 0.7 | -8.3 ± 0.0 | 3 |
| SL1-m | 885 ± 58 | 0.79 ± 0.07 | -6.7 ± 0.5 | -1.5 ± 0.5 | -8.2 ± 0.1 | 4 |
| SL1 | 73 ± 30 | 0.39 ± 0.05 | -5.8 ± 0.5 | -3.9 ± 0.7 | -9.7 ± 0.3 | 3 |
| SL1alt** | 2040 ± 359 | 0.68 ± 0.04 | -6.2 ± 0.5 | -1.5 ± 0.4 | -7.7 ± 0.1 | 3 |
| SL1-dII | 1220 ± 129 | 0.47 ± 0.05 | -3.5 ± 0.8 | -4.5 ± 0.9 | -8.0 ± 0.2 | 4 |
| SL1-p | 2210 ± 197 | 0.49 ± 0.03 | 6.6 ± 0.3 | -14 ± 0 | -7.6 ± 0.1 | 3 |
| ***A3m*** | | | | | | |
| SL1-p | 276 ± 17 | 0.34 ± 0.03 | 4.0 ± 0.4 | -13 ± 0 | -8.9 ± 0.0 | 3 |
| ***A12m*** | | | | | | |
| SL1-p | 396 ± 111 | 0.10 ± 0.01 | 9.3 ± 1.3 | -18 ± 1 | -8.7 ± 0.1 | 3 |
| ***ΔPYNT*** | | | | | | |
| SL1-p | 451 ± 262 | 0.11 ± 0.02 | 6.6 ± 0.2 | -15 ± 1 | -8.6 ± 0.3 | 3 |
| **Hexim1 dimer (syringe)** | | | | | | |
| SL1-dI | 458 ± 28 | 1.12 ± 0.05 | -5.8 ± 0.5 | -2.7 ± 0.5 | -8.6 ± 0.0 | 4 |
| SL1-m | 782 ± 26 | 0.87 ± 0.04 | -7.3 ± 0.3 | -0.9 ± 0.3 | -8.3 ± 0.0 | 3 |
| SL1 | 183 ± 18 | 1.75 ± 0.01 | -3.5 ± 0.0 | -5.5 ± 0.1 | -9.1 ± 0.1 | 3 |
| SL1alt** | 748 ± 15 | 1.01 ± 0.09 | -5.1 ± 0.0 | -3.1 ± 0.0 | -8.3 ± 0.0 | 4 |
| SL1-d(api) | 96 ± 22 | 1.40 ± 0.08 | -6.2 ± 0.2 | -3.3 ± 0.3 | -9.5 ± 0.2 | 4 |

* Number of individual experiments performed. Individual fits were performed for each experiment to calculate thermodynamic parameters and from these values, the average values and standard deviations were determined.
** Circular SL1alt in the ITC experiments refers to the UUCG tetraloop construct.

**Uncropped gel in Supplementary Fig. 4A**

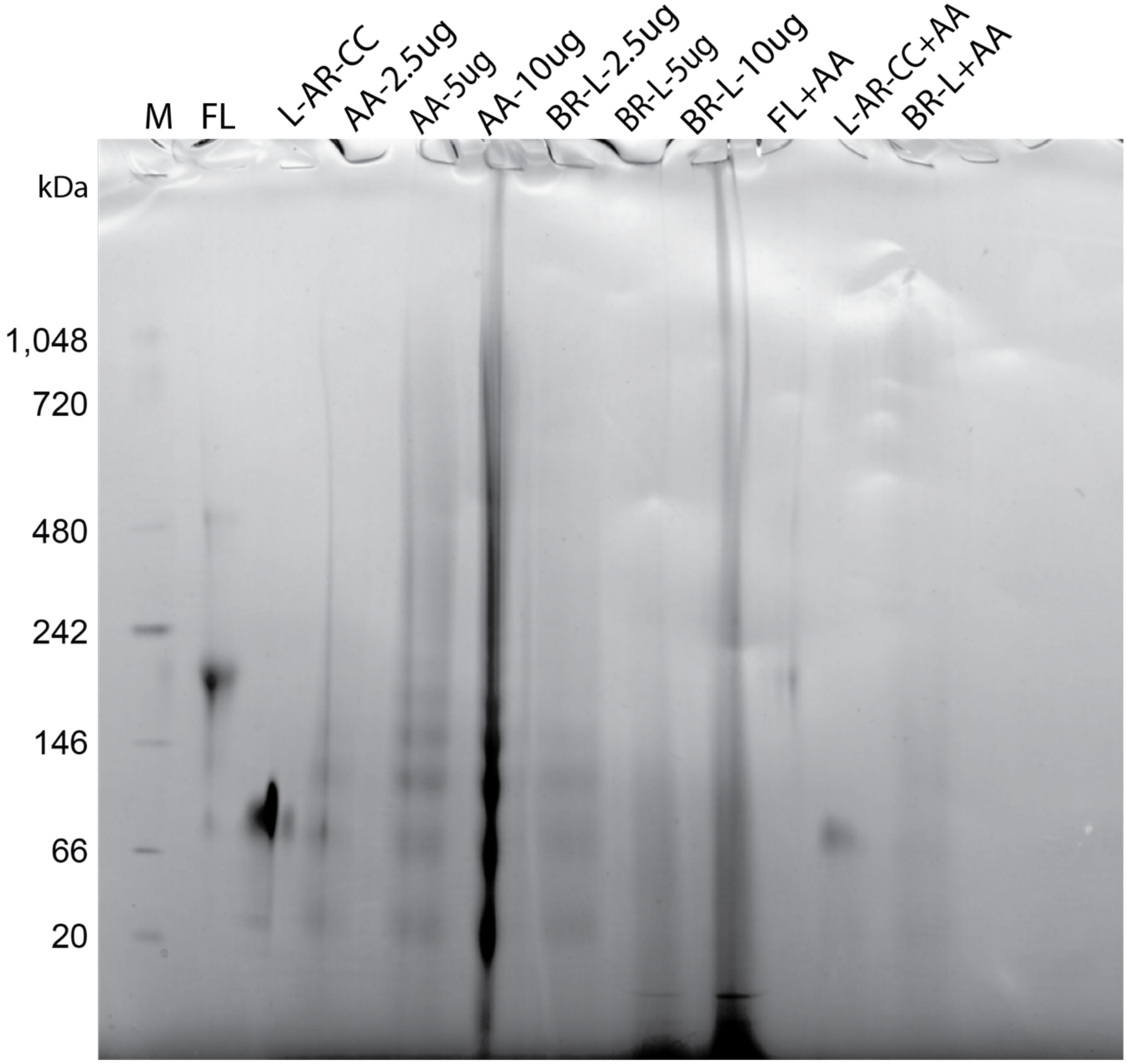

**Uncropped gels in Supplementary Fig. 14**

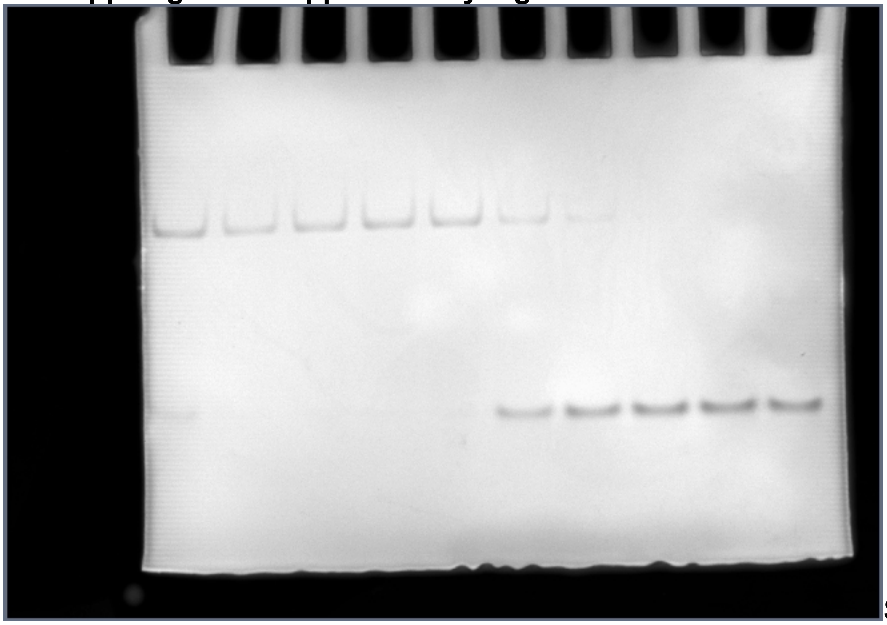

SL1-dI$_{\Delta U}$ + BR-L-AR

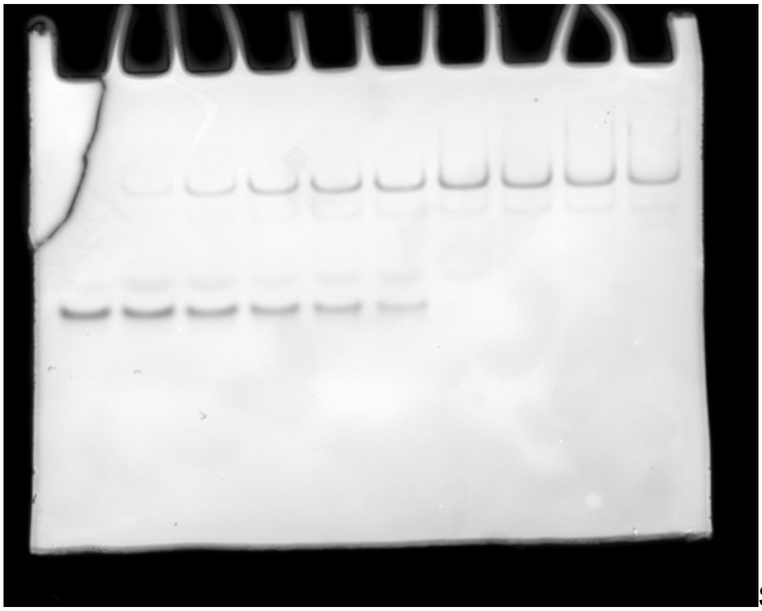

SL1-dI + BR-L-AR

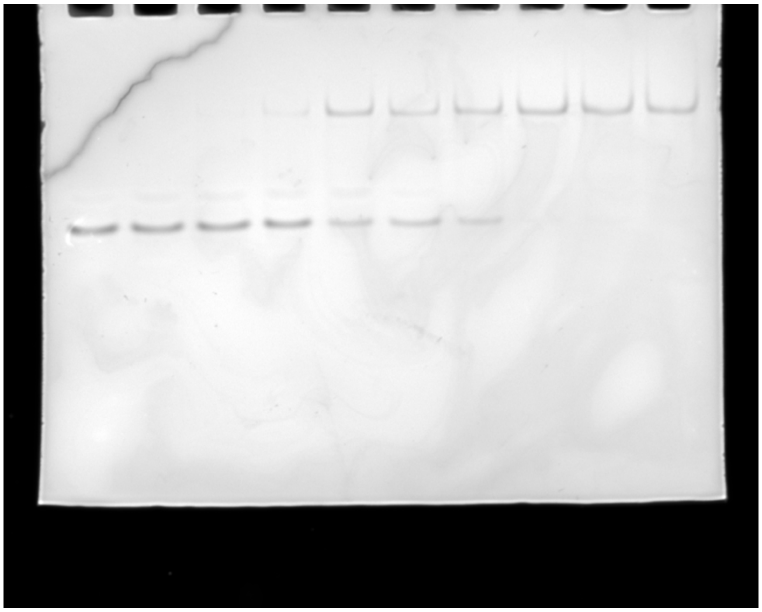

SL1-dII + BR-L-AR

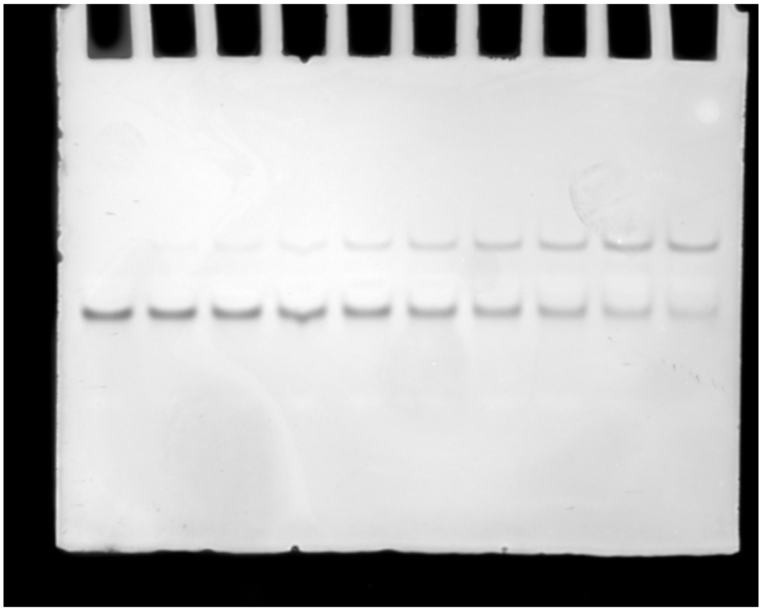

SL1-dI + BRext(149-179)

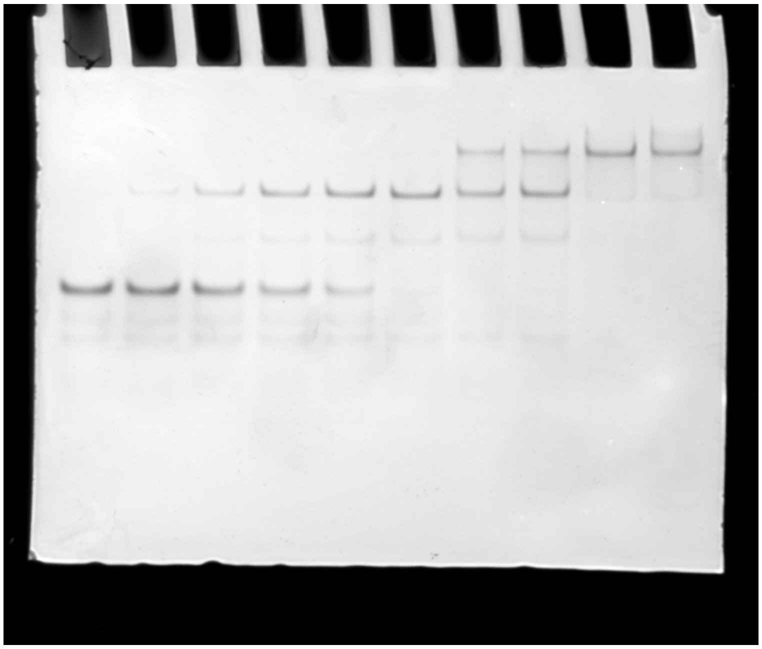

SL1-d$_{\Delta U}$ + BR-L-AR

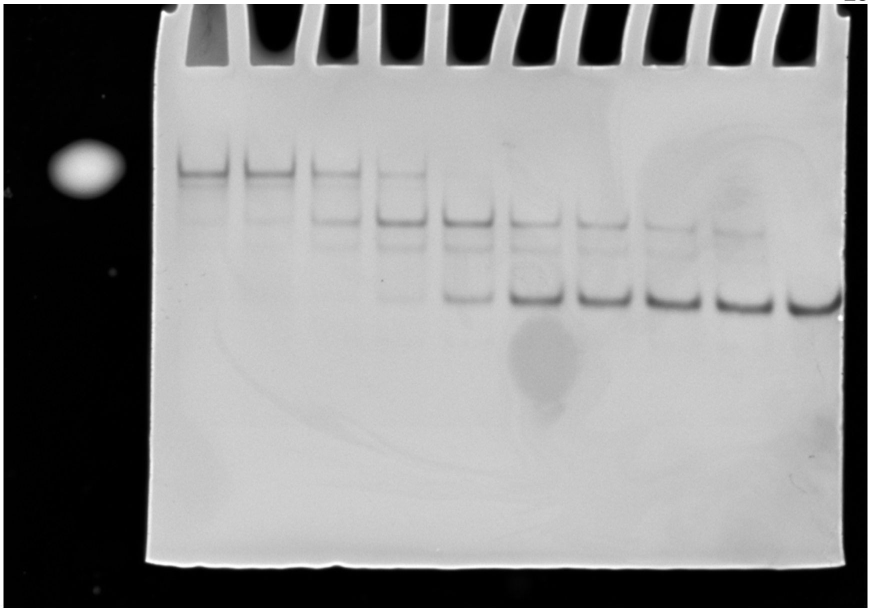

SL1-dI + BR-L(136-209)

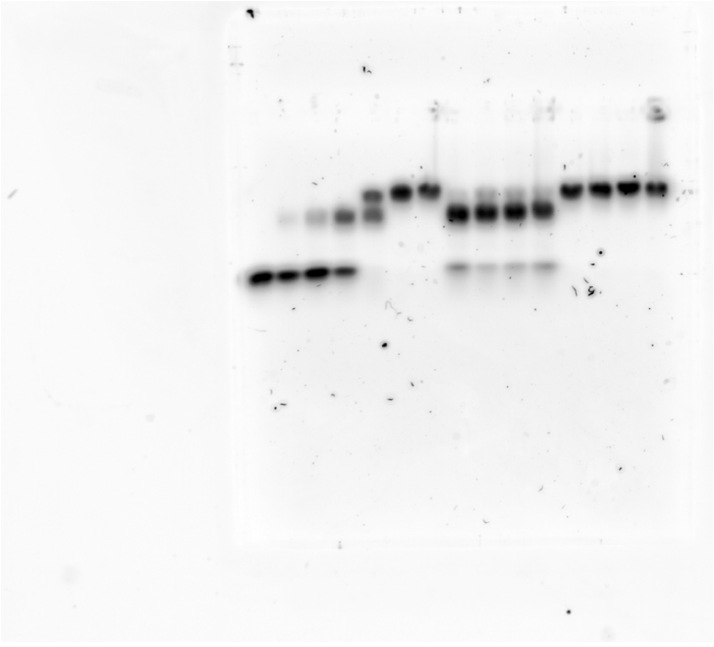

Linear SL1-SL4$^{ms2}$ + Hexim1

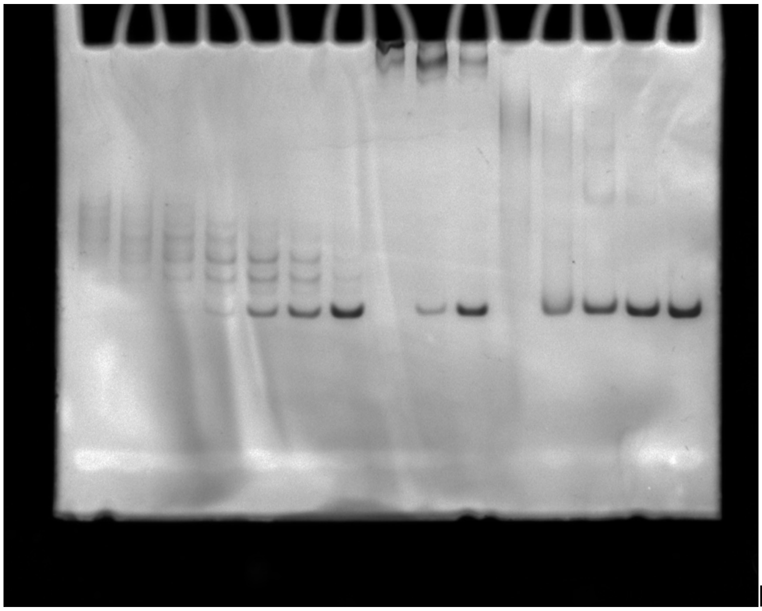

Linear SL1 + Hexim1 or BR-L(136-209)
